# Resource competition can explain simplicity in microbial community assembly

**DOI:** 10.1101/2023.06.13.544752

**Authors:** Hyunseok Lee, Blox Bloxham, Jeff Gore

## Abstract

Predicting the composition and diversity of communities is a central goal in ecology. While community assembly is considered hard to predict, laboratory microcosms often follow a simple assembly rule based on the outcome of pairwise competitions. This assembly rule predicts that a species that is excluded by another species in pairwise competition cannot survive in a multispecies community with that species. Despite the empirical success of this bottom-up prediction, its mechanistic origin has remained elusive. In this study, we elucidate how this simple pattern in community assembly can emerge from resource competition. Our geometric analysis of a consumer-resource model shows that trio community assembly is always predictable from pairwise outcomes when one species grows faster than another species on every resource. We also identify all possible trio assembly outcomes under three resources and find that only two outcomes violate the assembly rule. Simulations demonstrate that pairwise competitions accurately predict trio assembly with up to 100 resources and the assembly of larger communities containing up to twelve species. We then further demonstrate accurate quantitative prediction of community composition using harmonic mean of pairwise fractions. Finally, we show that cross-feeding between species does not decrease assembly rule prediction accuracy. Our findings highlight that simple community assembly can emerge even in ecosystems with complex underlying dynamics.

**Significance:** Multispecies microbial communities play an essential role in the health of ecosystems ranging from the ocean to the human gut. A major challenge in microbial ecology is to understand and predict which species can coexist within a community. While a simple empirical rule utilizing only pairwise outcomes successfully predicts multispecies laboratory communities, its mechanistic origin has remained unexplained. Here, we find that the observed simplicity can emerge from competition for resources. Using a generic consumer-resource model, we demonstrate that community assembly of highly complex ecosystems is nevertheless well predicted by pairwise competitions. Our results argue that community assembly can be surprisingly simple despite the potential complexity associated with competition and crossfeeding of many different resources by many different species.

## Introduction

Microbes coexist in complex, multi-species communities across scales. On a small scale, the microbiome affects health and disease of its hosts, including humans [1, 2, 3, 4, 5]. On a large scale, microbial communities play crucial roles for Earth’s biogeochemical cycles in oceans and soil [6, 7, 8, 9]. Understanding assembly of these communities (i.e. which species coexist and why) is a central goal in ecology that can impact agriculture, planetary science, and human health [10, 11, 12].

Predicting assembly is, however, a challenging problem: communities often have many species that interact in diverse ways, and characterizing all the potentially relevant interactions can be practically impossible [13, 14, 15, 16, 17, 18]. Similarly, theorists have found that community assembly prediction is an inherently difficult problem unless every information about the ecosystem is already given [19, 20, 21]. Even in simple models (e.g Lotka-Volterra model), predictions involve estimation of many parameters and may lead to over-fitting. Also, various factors such as higher-order, sublinear, and beyond-pairwise interactions, stochasticity, rapid evolution, and priority effects reinforce the unpredictability [22, 23, 24, 25, 26, 27, 28, 29, 30, 31]. In the end, since ecosystems are mostly too complex to be fully identified, both simple intuition and rigorous theory expect community assembly to be hard to predict.

Surprisingly, observed microbial community assembly is often predictable [32, 33, 34, 4, 35, 36, 37, 38]. In particular, the assembly outcome in laboratory microcosms from an initially diverse species pool can be determined bottom-up from pairwise competitions among the species without any model assumption nor parameter fits^1^. Friedman et al 2017 formulated this purely empirical bottom-up prediction as the ‘assembly rule,’ proposing that species A will exclude species B in community assembly if A excludes B in pairwise competition [34] (Fig. 1A) ^2^. To test this idea, they performed serial dilution experiments and found that the assembly rule predicted species survivals in trio communities with high accuracy (89.5%). This assembly rule has since also predicted community assembly in the intestine of the worm *C. elegans* and amidst changes in the environment due to variation in the dilution rate and temperature [34, 39, 40, 41, 42]. The rule’s frequent experimental success highlights how, despite the expected complexity, community assembly can be predicted in a surprisingly simple way. Yet the condition for simplicity remains unclear because the assembly rule, being purely empirical, does not provide deeper explanations for why it works nor when to expect violations. A possible mechanistic origin for the empirical assembly rule is a crucial missing piece toward a better understanding of community assembly and its predictability.

**Fig 1.**
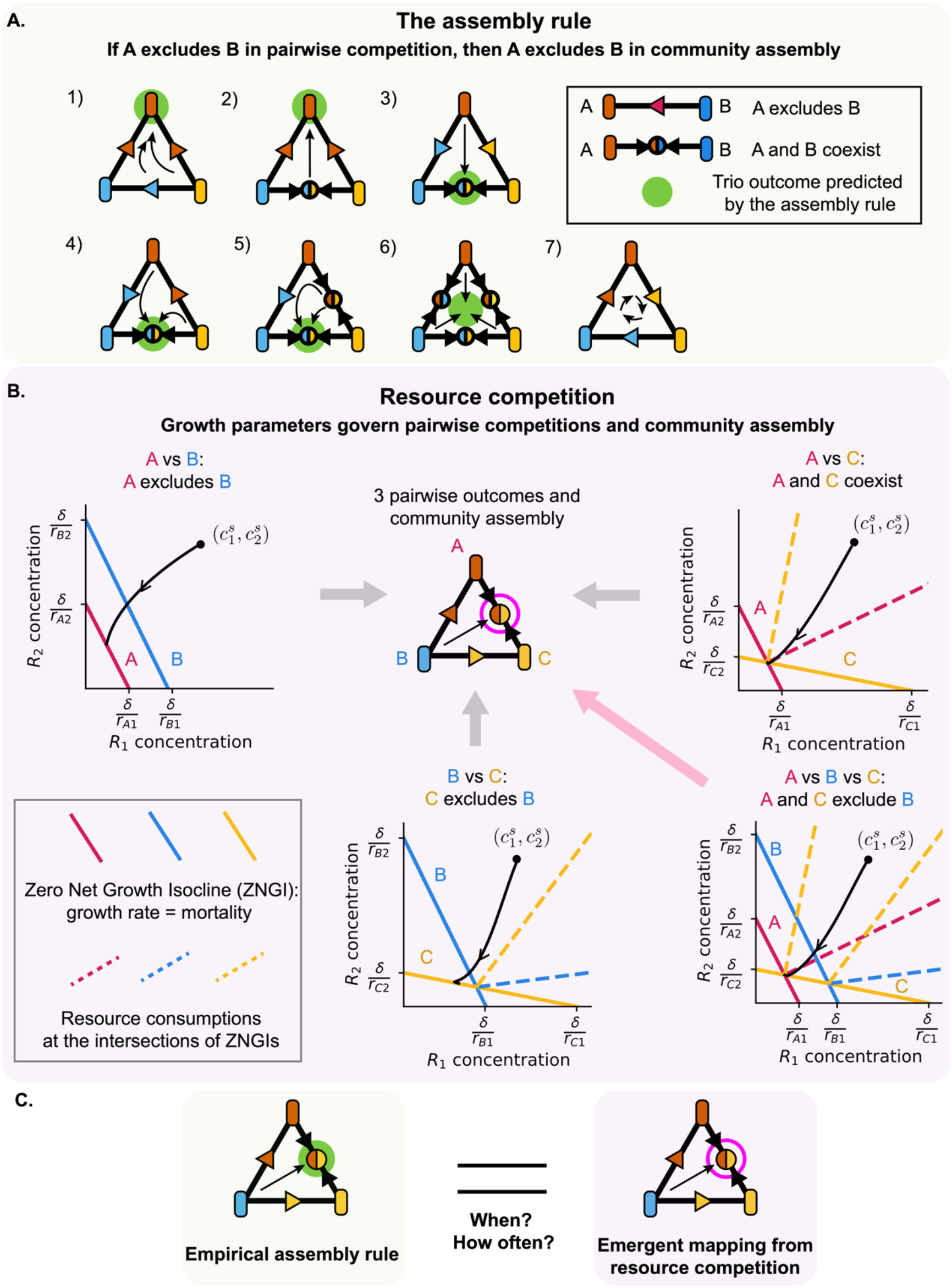
Assembly rule may emerge from resource competitions. **a**. Assembly rule predicts trio competition outcomes based on pairwise competition outcomes. The assembly rule predicts that if species A excludes species B in pairwise competition, then A excludes B in community assembly. Equivalently, the assembly rule assumes that a species invades a community if and only if it can invade every member of the community. The rule predicts exactly one trio outcome for each combination of pairwise competitions except for the rock-paper-scissors case for which no trio outcome is predicted. **b**. In resource competition models, growth parameters govern both pairwise competitions and community assembly. Each competition is represented in the space of resource concentrations using ZNGIs (Zero Net Growth Isocline: the set of resource concentrations on which a species’ growth rate equals its mortality rate and thereby the species maintains a nonzero population size) and resource consumptions at the intersections of ZNGIs (i.e. lines from each intersection 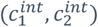 in the direction of 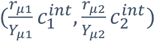. Each resource consumption vector represents a species’ resource consumption when the resource concentration of the environment is at the intersection of ZNGIs, and this serves as the phase boundary between pairwise coexistence and exclusion (also see Fig. S1). The black dots represent resource supply 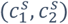 and the black curves represent trajectories of resource concentrations until the population reaches an equilibrium. Under this graphical representation, equilibrium concentrations must lie on the innermost ZNGIs and resource consumption must balance with resource supply at equilibrium. Followingly, for the illustrated example, A excludes B (top left), A and C coexist (top right), C excludes B (bottom center), and in trio competition A and C exclude B (bottom right). Finally, all the three pairwise outcomes and community assembly are represented on a simplex (top center). Geometric analysis of each competition is detailed in Results. Notably, this trio community follows the assembly rule. **c**. The emergent mapping from resource competition in Fig. 1B agrees with the assembly rule. Elucidation on when and how often this agreement holds may explain the origin of simple empirical rule’s successful predictions.

Competition for resources is a ubiquitous mechanism that often drives microbial community dynamics; for example, many studies have revealed how community composition depends on resource composition supplied to the environment [43, 44, 45, 38, 46, 47]. In models of resource competition, the interspecies interactions are mediated by the uptake (and potential release) of resources, introducing potentially complex dynamics and the emergence of “higher-order” interactions between species [28, 27, 48]. However, the same growth parameters govern both pairwise competitions and multi-species competitions (Fig. 1B). This implies the possibility of an emergent mapping – one that is not explicitly included in the model definition but instead arises from the community dynamics – between their outcomes. In this way, a bottom-up prediction from pairwise competitions to larger community assembly may be formulated under the framework of resource competition. Elucidating this bottom-up correspondence would shed light on the unexplained success of simple rule in predicting community assembly (Fig. 1C).

Here, we demonstrate that the emergent mapping between pairwise competitions and community assembly under resource competition nearly always agree with the empirical assembly rule. We first analyze a resource competition model with geometric principles and find that trio assembly always follows the assembly rule unless every species grows faster than a competitor on some resource. Moreover, we identify all possible combinations of pairwise and trio outcomes under resource competition for three resources and show that only two combinations violate the assembly rule. To verify this theoretical analysis and extend it to larger number of resources, we perform simulations of trio communities with up to 100 resources. Regardless of the number of resources, simulations of resource competition agree with the assembly rule prediction of species survival with over 90% accuracy and the list of outcomes agree with our geometric analysis.

Furthermore, simulations with larger communities and with cross-feeding between species continue to be predictable by the assembly rule. Our results highlight that a complex system with many microscopic parameters can nevertheless lead to predictable community assembly.

## Results

### Resource model leads to a simple mapping from pairwise outcomes to community assembly

To study community assembly under resource competition, we focus on a chemostat-like linear resource consumption model, which is a variation of MacArthur’s consumer-resource model with an external resource supply and universal dilution rate [49, 50, 51, 46] (Methods). The model assumes that per-capita growth rate is the sum of growth on each resource, which is proportional to concentration of the resource. This implies the resources are substitutable. While this choice of model simplifies some analysis, our main results hold multiple for other forms of resource competition models as well (SI Appendix I.1). The model dynamics are:

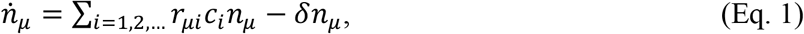

and

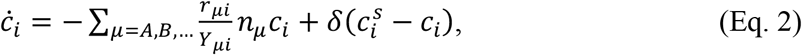

where *n*_*μ*_ is the population density of species *μ, c*_*i*_ is the concentration of resource *i* in the media, *r*_*μi*_ is the contribution of resource *i* to the per-capita growth rate of species *μ, Y*_*μi*_ is the biomass yield of species *μ* on resource *i, δ* is the dilution rate in chemostat which controls both universal mortality and resource supply rate, and 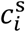 is the external supply concentration of resource *i*. In this paper, we will assume that relative biomass yields between resources are the same for all species (i.e. *Y*_*μi*_ =*a*_*μ*_*y*_*i*_). Under this assumption, each competition converges to a single equilibrium independent of initial population densities (i.e. multi-stability is not possible). We also explore the cases with variable yields and density-dependent outcomes in SI Appendix V.

Before we move on to our original results, here we briefly discuss our tool for studying the model: a graphical approach in the space of resource concentrations [52, 53, 54]. This approach utilizes each species’ Zero Net Growth Isocline (ZNGI), which is the set of resource concentrations for which growth of a species exactly keeps up with the dilution rate (Fig. 2) [54]. Notably, the equilibrium resource concentration satisfies two geometric principles. First, equilibrium concentrations must lie on the innermost ZNGIs such that all species’ populations are either constant or going extinct. A species survives if the equilibrium resource concentration is on its ZNGI [54]. Second, for resource consumption to balance supply, the supply must fall within the convex hull of the surviving species’ resource consumptions at the equilibrium [55]. Geometrically, this can be determined by drawing resource consumption vectors originating from all ZNGI intersections (Fig 1B) [51]. The first principle dictates whether species can ever coexist, and the second principle tells whether they coexist under a particular resource supply.

**Figure 2.**
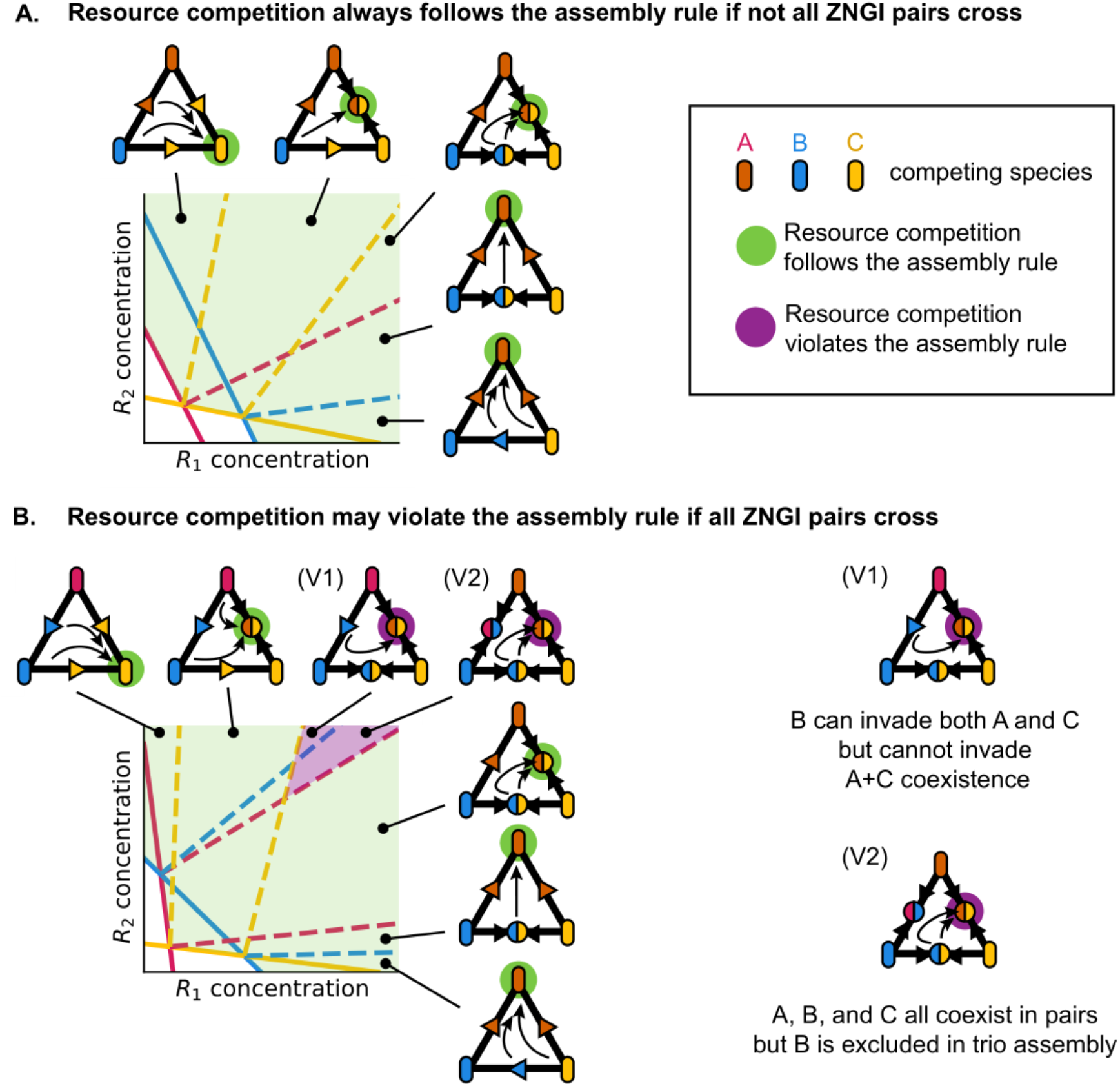
Geometric analysis of resource competition model reveals the conditions for assembly rule agreement and violation. **a**. Resource competition of trio community always follows the assembly rule if not all ZNGI pairs cross. In the illustrated example, species A’s ZNGI is inside species B’s ZNGI. Then A always outgrows B under any resource concentration, and thus B is excluded by A in both pairwise competition and community assembly. As a result, the community assembly follows the pairwise competition between A and C, and the assembly rule is always obeyed. In the illustrated example, while competition outcomes shift as the resource supply changes, each combination of pairwise and trio outcomes under any resource supply follows the assembly rule (Fig. S2). **b**. Resource competition of trio community may violate the assembly rule if all ZNGI pairs cross. In the illustrated example, each pair’s ZNGIs intersect. Then any species can outcompete its competitor at some resource concentration, and a third species may shift the competition outcome between a pair by modifying the resource environment. The illustrated example exhibits two violations (Fig. S2). In the violation V1 (with two pairwise coexistence and an exclusion), A and C survive in the community assembly while the assembly rule expects B to survive instead of A. By coexisting with C, A maintains the resource concentration on which it outgrows B, which results in the unexpected exclusion of B in community. In the violation V2 (with three pairwise coexistence), only A and C survive while the assembly rule expects all A, B, and C to survive. As in the previous case, by coexisting, both A and C maintain the resource concentration on which they outgrow B, resulting in the unexpected exclusion of B. In a generic resource competition, all-species coexistence requires all-pair coexistence but not vice versa (Fig. S3).

Fig. 1B illustrates these geometric principles for pairwise and trio competitions. Between species A and species B, A’s ZNGI is always inside B’s ZNGI, so, by the first geometric principle, if only A and B are present any fixed point will have A surviving and B being driven extinct. By contrast, both A and C’s ZNGIs and B and C’s ZNGIs intersect, so the first principle suggests both these pairwise competitions may lead to coexistence. However, the second geometric principle now becomes relevant: the resource supply lies between A and C’s consumption but not between B and C’s consumption. Thus, while A and C coexist, B and C cannot because no population composition can balance the resource supply and keep the resource concentrations at the intersection of B and C’s ZNGIs. In the trio competition, the same two principles apply: B cannot survive because its ZNGI is never one of the innermost while A and C again coexist because the resource supply lies between their resource consumption. This trio outcome also matches the assembly rule. While the graphical analysis of each competition here is not new, the relationship between pairwise competitions and community assembly have not been elucidated with this approach.

The graphical approach identifies conditions for resource competition to follow the assembly rule. Remarkably, for trio communities, resource competition outcomes always match with the assembly rule predictions when not all the ZNGIs cross. When species A’s ZNGI is always below species B’s ZNGI, then species B will be excluded in both pairwise competition between A and B and trio community assembly. This implies that the community assembly always follows pairwise competition between A and C. Thus, under a generic resource competition model (e.g. with non-linear ZNGIs), existence of a non-intersecting ZNGI pair is a sufficient condition for the assembly rule to be obeyed in a trio community assembly. Fig. 2A illustrates this agreement: while both pairwise and trio competition outcomes shift depending on the resource supply, every set of pairwise and trio outcomes follow the assembly rule.

Resource competition outcomes can violate the assembly rule when all ZNGI pairs cross. In Fig. 2B, while the assembly rule is obeyed for most resource supply concentrations, a few violations are permitted. In the violation V1 (with two coexisting pairs and one excluding pair), species A and C survive while the assembly rule expects B to survive instead of A. While B excludes A in pairwise competition, because A’s and B’s ZNGIs cross, A could exclude B under a different resource supply. By coexisting with C, A maintains the resource concentration on which it outgrows B, resulting in an unexpected trio outcome. Similarly, in the other violation V2 (with three coexisting pairs), species A and C survive while the assembly rule expects A, B, and C to all coexist. As in the previous case, by coexisting, A and C both maintain the resource concentration on which they outgrow B, resulting in the unexpected exclusion of B. In fact, when all pairs coexist in a two-resource environment their community assembly always results in a two-species survival, unless parameters are fine-tuned to have a mutual intersection of all ZNGIs [55]. More generally, under a generic resource competition model, all-species coexistence requires all-pair coexistence but not vice versa (Fig. S3). Overall, how these violations can occur highlight that resource competition inherently involves higher-order interactions; the existence of a third species can modify the resource concentrations in the environment, and this leads to a change in the resource-mediated interaction between a pair [28, 56].

So far, we have investigated how resource competitions among some trios may follow or violate the assembly rule depending on the resource supply. Extending this procedure to all possible trios and resource supplies would identify the complete list of resource competition outcomes (Fig. 3A). While this may sound formidable, there are only a few topologically distinct ways that 3 ZNGIs can intersect. By enumerating all the possible phase diagrams, we identify all pairwise and trio outcomes under two and three resources (Fig. 3B, SI Appendix II & III). Interestingly, the only possible violations of the assembly rule are the two previously identified cases (Fig. 2B). All other resource competition outcomes follow the assembly rule. The list of possible resource competition outcomes has several notable features (Fig. 3C). First, resource competition can realize every outcome predicted by the assembly rule. Second, while resource competition can violate the assembly rule, both violations have been experimentally observed [34]. Strikingly, one of the two violations in the model (two-species survival under all-pair coexistence) is the most frequently observed violation in experiments [34]. Third, resource competition excludes many possible assembly rule violations, including the violations allowed under the Lotka-Volterra model [57]. In addition, rock-paper-scissors pairwise outcome, for which assembly rule prediction does not exist, is impossible in our resource competition model. In the end, the overall agreement between resource competition and the assembly rule prediction suggests that resource competition often results in simple community assembly.

**Figure 3.**
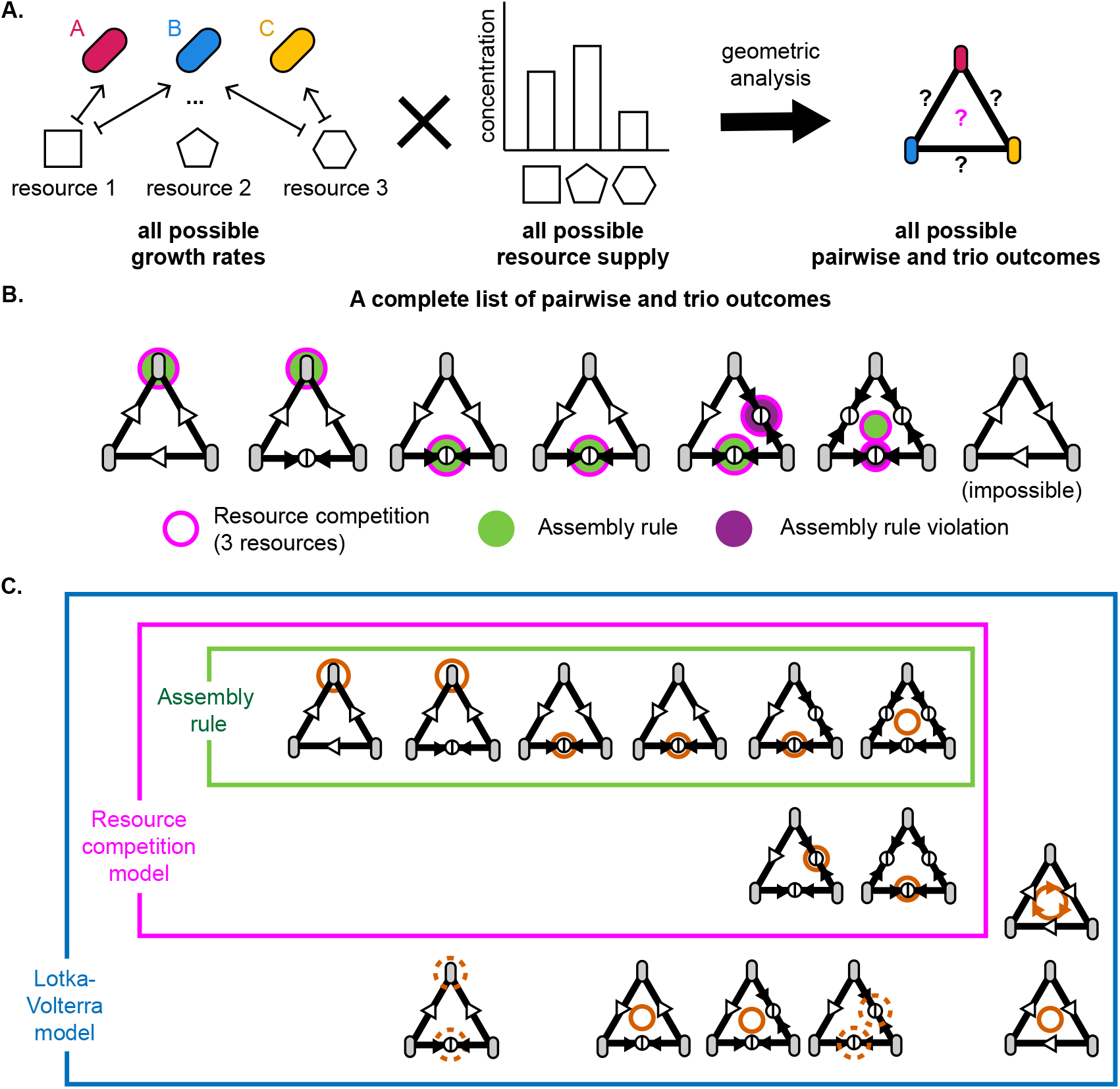
A complete list of pairwise and trio outcomes under resource competition for three resources. **a**. We find all possible pairwise and trio outcomes under resource competition by investigating outcomes under all possible growth rates and resource supply. Geometric analysis simplifies this procedure because there are only a few distinct arrangements of ZNGIs under 1, 2, and 3 resources. **b**. A complete list of pairwise and trio outcomes under resource competition for three resources (SI Appendix II & III). **c**. Trio outcomes under the assembly rule, resource competition model, and Lotka-Volterra model. Dotted circles imply bistable community assembly. Resource competition model can lead to all assembly rule predictions plus only two assembly rule violations. Rock-paper-scissors pairwise outcome, for which assembly rule prediction does not exist, is impossible in our resource competition model.

Before proceeding, we note that not all the observed violations of assembly rule are allowed in the resource competition model we considered [34, 58]. A particularly interesting case is the all-species coexistence in community assembly accompanied by one or more pairwise exclusions (Fig. S3). This violation thus can be sign of another mode of interaction (e.g. Lotka-Volterra model allows this violation), fluctuating environment, community under oscillation or chaos instead of a single fixed point, species directly modifying other competitors’ traits, and many other possible ways to break our model assumptions.

### Resource competition in complex ecosystems often follows the assembly rule

To augment the geometric analysis of resource competition under two and three resources, we simulated resource competition across different numbers of supplied resources from 1 to 100 (Fig. 4A, Methods). We compared the simulated community assembly with three different predictions: ‘Everyone survives’ that always predicts all three species to survive, ‘Fastest grower survives’ that predicts the best grower under the resource supply to exclude other two species, and the assembly rule (Methods).

**Figure 4.**
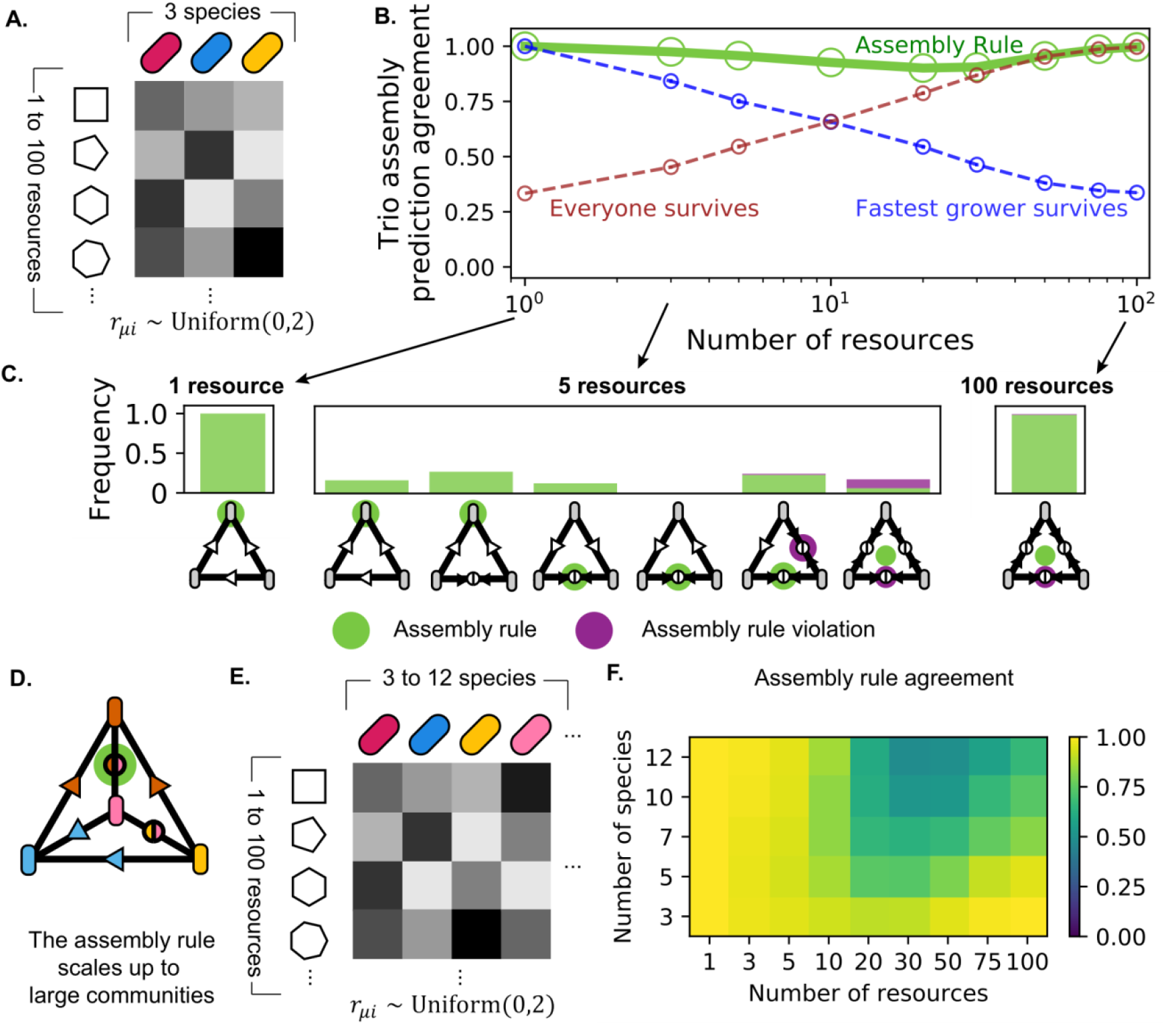
Simulations of resource competition typically followed the assembly rule. **a**. We simulated resource competition of trio community under variable number of resources from 1 to 100. The growth rates were sampled from a discretized uniform distribution from 0 to 2 (Methods). In the SI, we show that the results here hold also when a metabolic tradeoff constrains growth rates, species do not utilize all resources, growth rates are not uniformly distributed, or growth rates are closer to dilution rates (SI Appendix I.2, I.3, and I.4.). **b**. In simulations, community assembly often followed the assembly rule. We compared the simulated resource competitions with three different predictions: in addition to the assembly rule, ‘Everyone survives’ always predicts all three species’ survival, and ‘Fastest grower survives’ predicts the best grower under the resource supply to exclude other two species. We quantified prediction agreements by fraction of species in each trio that matched each prediction. Overall, across all numbers of supplied resources, the agreement between resource competition and the simple assembly rule was maintained above 90%. ‘Fastest grower survives’ prediction failed in many-resource regime and ‘Everyone survives’ prediction failed in few-resource regime. **c**. Distributions of simulated outcomes under 1, 5, and 100 supplied resources. The bar plots show observed frequency of each set of pairwise outcomes and community assembly. Community assembly that followed the assembly rule is marked in green, and violations are marked in violet. With single supplied resource, fastest grower on the single resource excluded others in all competitions. With 5 supplied resources, the outcomes became diverse and the assembly rule was no longer perfect. Nevertheless, simulations agreed with geometric analysis; all expected outcomes including the two violations were observed, and no unexpected outcome was observed (Fig. 3B). With 100 supplied resources, due to the large number of available niches, species almost always coexisted in both pairwise and trio competitions. Simulations with other numbers of supplied resources showed the same trend (Fig. S4). **d**. Assembly rule scales up to large communities. For example, in the illustrated 4-species competition, since species B and C are both excluded by A in pairwise competitions, they are also excluded in 4-species competition. **e**. We simulated resource competition of variable number of species (from 3 to 12) under variable number of resources (from 1 to 100). The growth rates were sampled from a discretized uniform distribution from 0 to 2 (see Methods) **f**. The assembly rule works best with few species or few resources. Resource competitions of multispecies communities often followed the assembly rule when a small number of resources were available. Similarly, resource competitions of communities with small number of species often followed the assembly rule even when many resources were available. With large number of both species and resources, assembly rule no longer worked well.

To quantify the agreement between the assembly rule and resource competition, we calculated the accuracy with which the assembly rule predicted whether each individual species survived (Fig. 4B). Remarkably, the average agreement between the assembly rule and resource competition never dropped below 90% (minimum at 10 resources, with accuracy 90.13% ±0.01%, SEM, N=500). This species-by-species quantification was chosen for consistency with previous work and for being a simple metric that can be applied to communities of many species (see below) without indicating a complete failure when only a single species is inaccurately predicted [34]. This uniformly high accuracy of the assembly rule prediction highlights the simplicity and predictability of resource-competing communities. In the SI, we show that the results here hold also when a metabolic tradeoff constrains growth rates, species do not utilize all resources, growth rates are not uniformly distributed, or growth rates are closer to dilution rates (SI Appendix I.2, I.3, and I.4.).

When only one resource was supplied, the fastest grower excluded all other species, and the assembly rule was always successful (Fig. 4C). In the other limit, where 100 resources were supplied, the number of available niches became much greater than the number of species. Therefore, species mostly survived in all pairwise and trio competitions, and the assembly rule was again successful. In the end, the assembly rule worked almost perfectly on both few-resource and many-resource limits. With an intermediate number of supplied resources, resource competitions no longer perfectly agreed with the assembly rule. However, any simulated violation was one of the two outcomes expected from our geometric analysis, suggesting that increasing the number of supplied resources did not enable new unexpected outcomes (Fig. S3). Similarly, violations never occurred when some ZNGI pairs did not cross (Table S1), again confirming that insights gained from our geometric analysis remained accurate and valuable in more complex environments.

In addition to competition for many resources, we also considered another axis of complexity: competition among more than three species. We note that the assembly rule is applicable to communities larger than trio (Fig. 5A). We simulated community assembly among 3, 5, 7, 10, and 12 species along with the corresponding pairwise competitions (Fig. 5B). For a fixed number of species, the non-monotonic trend of assembly rule prediction accuracy reached minimum at intermediate number of resources (Fig. 5C). And as the number of species increased, the minimum shifted toward larger number of resources and lower prediction accuracy. This shows that the assembly rule prediction often works for small communities with any number of resources, as observed experimentally [34]. Similarly, the assembly rule predicted community assembly well when there are few available resources. For communities with large numbers of both species and resources, though, assembly rule prediction no longer worked well. This result may guide us when to use the assembly rule in practice.

**Figure 5.**
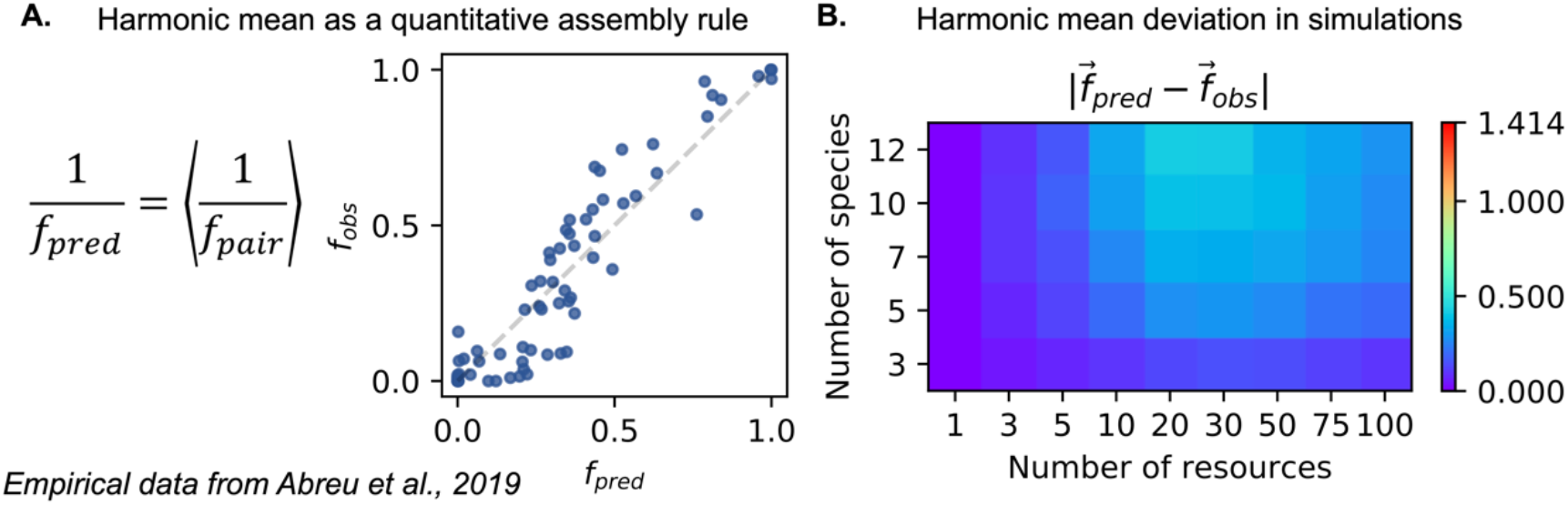
Community composition often follows harmonic mean of pairwise competition outcomes. **a**. Harmonic mean of pairwise fractions quantitatively predicts experimental microcosm. While the assembly rule is simple and useful, a quantitative prediction of community composition can provide additional fine-grained information. We found that, in the limit of few species and resource specialization, community composition can be approximated with harmonic mean of pairwise fractions (SI Appendix IV). We tested this harmonic mean prediction with empirical data from an experimental study [39]. The harmonic mean predictions from pairwise outcomes were close to observed community compositions (L2-distance of 0.17 ± 0.02, SEM, N=23), with the mean difference between prediction and observed fractions being 0.09 ± 0.01, SEM, N=69. **b**. Harmonic mean of pairwise fractions predicts simulated community compositions. We tested the harmonic mean prediction with simulations of resource-competing communities (Fig. 4D & 4E). With L2-norm distances, we find that the prediction error becomes largest at intermediate number of resources and large number of species. Nevertheless, compared to the maximum possible L2-norm distance = √2, the prediction error stays small even in complex communities.

### Harmonic mean as a quantitative community assembly rule

So far, we have used the assembly rule to predict whether a species would survive or go extinct in community assemblies. While such binary implementation is simple and useful, it also raises the question of whether a quantitative prediction of community assembly is possible.

We found that, under our resource competition model, it is possible to calculate exact species fractions from pairwise outcomes when each species consumes only one resource (SI Appendix IV). In the limit of few surviving species, this result can be approximated by the harmonic mean of fractions in pairwise competitions:

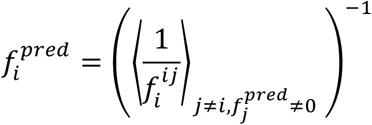

where 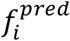 is the predicted fraction of species *i*, 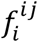 is the fraction of species *i* in a pairwise competition between species *i* and *j*, and ⟨… ⟩ is a simple average. The calculation is done iteratively to consider pairwise competitions between surviving species only (e.g. if A excludes B and B excludes C, then C survives in trio assembly if it coexists in with A in pairwise competition). Note that this quantitative prediction recapitulates the assembly rule’s definition that A excludes B in community assembly if A excludes B in pairwise competition, i.e. any 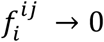 would lead to 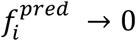.

To test this harmonic mean prediction, we applied it to data from an experimental study [39]. The harmonic mean successfully predicted species fractions in experimental community assemblies (Fig. 5A). We also quantified overall prediction accuracy with L2-norm distance between predicted and observed community compositions, 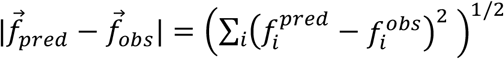. The harmonic mean predictions from pairwise outcomes were close to observed community compositions (0.17 ±0.02, SEM, N=23), with the mean difference between prediction and observed fractions being 0.09 ±0.01, SEM, N=69.

Next, we tested the harmonic mean predictions with simulated resource-competing communities. Individual species fractions were well predicted in cases of few resources or few species, and yet underestimated fractions of surviving species when both species and resources were in large numbers (SI Appendix IV). Interestingly, the systematic deviation between the observed and predicted fractions is similar to that observed in the experiment (SI Appendix IV, Fig. 5A).

Similarly, with L2-norm distances, we find that the prediction error becomes largest at intermediate number of resources and large number of species (Fig. 5B). For example, the mean prediction error was 0.12 ±0.01, SEM, N=500 for 5 species and 5 resources and 0.42 ±0.01, SEM, N=100 for 12 species and 20 resources. Nevertheless, compared to the maximum possible L2-norm distance = √2, the prediction error stays small even in complex communities.

### Community assembly in the presence of cross-feeding typically follows the assembly rule

So far, we have assumed that species always consume resources and thereby all interactions are antagonistic. In nature, however, microbes often produce metabolic byproducts that are consumed by another species. This cross-feeding increases the number of available niches and increases the number of species that could potentially coexist in community [59, 49, 43, 45, 60]. To test whether simplicity and predictability in assembly are maintained under the presence of cross-feeding, we implemented conversions from one resource to another that are proportional to consumption of the first resource (Fig. 6A, Methods). We also assumed hierarchy in resource complexity such the most complex resource may be converted into any of less complex resources and the least complex resource may never produce another metabolite. Assuming fully connected network without any hierarchy did not change the result (SI Appendix I.5). We also assumed that the cross-feeding reduces the efficacy of the first resource as its energy content is leaked out [45]. The link between each resource pair (from more complex resource to a simpler one) was established with probability *p*, and *l* is the conversion rate from consumption of a complex resource to production of a simpler “metabolite” resource (Fig. 6B, Methods). Fig. 6C and 6D show the simulation results with three species, single supplied resource, and 9 potential metabolites under varying sparsity and strength of cross-feeding. We also simulated cross-feeding communities with varying number of species and resources and found no significant changes in assembly rule prediction accuracy (SI Appendix I.5).

**Figure 6.**
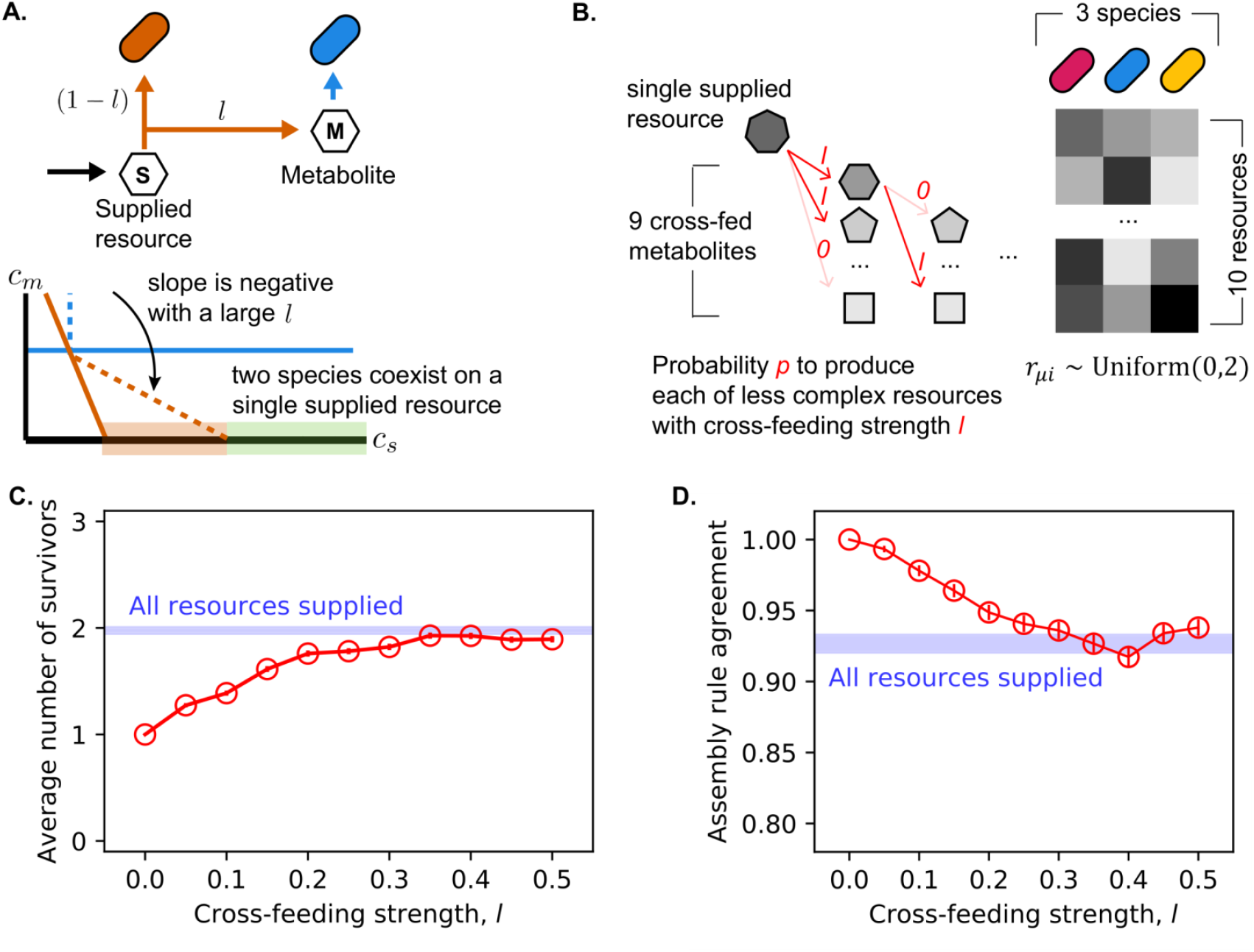
Resource competition with cross-feeding is often simple. **a**. Cross-feeding enables coexistence of two species under single supplied resource. As a species consumes a resource, fraction l of the resource can be leaked out as metabolic byproducts which others can consume. This increases the number of niches in the environment and allows for more species to coexist. **b**. We ran simulations of trio competing under cross-feeding. Only the most complex resource was supplied, but cross-feeding could allow coexistence as 9 other resources could appear as metabolites. Each trio shared a cross-feeding network in which each resource may cross-feed simpler resources with probability p and cross-feeding strength l (see Methods). For simplicity, we fixed p=0.5 and varied l in simulations. **c**. Number of survivors in trio competitions increased with cross-feeding strength. With l=0, only one species survived in trio competition as expected. The number of survivors could reach, but could not exceed, the number of survivors with all 10 resources being externally supplied instead of being cross-fed. **d**. Prediction accuracy of the assembly rule was still high for cross-feeding communities. The prediction accuracy could approach, but could not be significantly lower than, the prediction accuracy with all 10 resources being supplied.

As cross-feeding became stronger, the number of species surviving in community assembly tended to increase due to the creation of additional niches (Fig. 6C). Compared to the case where all 10 resources are externally supplied without any cross-feeding (average number of survivors = 1.98), a cross-feeding community could reach the same biodiversity with sufficiently large cross-feeding strength *l*. Also, as *l* increased, the assembly rule agreement of communities with cross-feeding approached to the case where all 10 resources were externally supplied (Fig. 6D). Overall, assembly rule prediction accuracy stayed above 90% (minimum at *l=0*.*4*, prediction accuracy 91.73% ±0.01%, SEM, N=500). Thus, the simplicity of community assembly under resource competition was maintained even when cross-feeding complicated the underlying dynamics.

## Discussion

In this study, we demonstrated that competition for resources may lead to the experimentally observed simplicity in community assembly. Trio assembly is always predictable from pairwise competitions if a species grows better than another species on every supplied resource.

Otherwise, resource competition may violate the assembly rule, but only via two rare and specific scenarios. This predictability holds for communities with large number of species and resources and in the presence of cross-feeding.

The simplicity that emerges from our resource competition model is at odds with the model’s complexity. Compared to the Lotka-Volterra model with 6 pairwise interactions for a trio, our resource competition model for multiple resources has more parameters and allows for higher-order interactions in which a third species affects pairwise competition between others. Yet resource competition leads to a more constrained set of outcomes than the Lotka-Volterra model. This is because, in the resource competition model, species modify the environment but not traits of competitors, e.g. each species’ ZNGI stays the same regardless of whom it competes against. This invariance of the traits of individual species, such as per-capita per-resource growth rates, over different competitions enables a simple emergent mapping despite the underlying complexity.

Geometric analysis provides insights on the simple community assembly. The relative ordering between ZNGIs determines a competitive hierarchy between species, and simple arrangement of ZNGIs leads to simple community assembly. Specifically, when a ZNGI does not cross with another ZNGI, then the trio community always follows the assembly rule. This competitive hierarchy also explains why resource competitions are often transitive and rock-paper-scissors pairwise outcome is impossible. In addition, geometric principles impose hierarchy between coexistences such that coexistence of many species require coexistence of all pairs between them: A, B, and C coexist as a trio community only if A-B, A-C, and B-C pairs all coexist, but not vice versa^3^. These geometric hierarchies provide a way to understand the agreement between bottom-up assembly rule prediction and resource competition.

Like in laboratory experiments, community assembly under resource competition sometimes deviates from the bottom-up assembly rule prediction. Strikingly, such violations may strengthen the connection between experimentally observed community assembly and resource competition. For example, we find that the most frequently observed violation in the experiment is the most frequently observed violation in simulations [34]. Our resource model analysis also clarifies conditions for the bottom-up prediction’s failure in terms of growth rates and resource supply. Trio community assembly may violate the assembly rule only when all three ZNGI pairs cross. This also implies that metabolic tradeoffs may lead to frequent violations of the assembly rule, and indeed some cases of metabolic tradeoff result in more violations in our simulation (SI Appendix I.2). Relatedly, experiments have found that co-evolution or history of coexistence may lead to more frequent violations [42, 58]. Beyond trio communities, assembly rule failure increases with the number of species, especially when dozens of resources are supplied (Fig. 4). Experiments aided with metabolomics may verify the criteria and further improve the predictive power [61].

While we focused on a simple linear consumption model, there are many other models for ecological interactions [62]. For example, the Lotka-Volterra model allows more community assembly outcomes from the pairwise competitions compared to our resource competition model. While our resource competition model and the assembly rule successfully describe laboratory microcosms, further work will be necessary to determine the rules of community assembly in other systems that may involve various interactions such as toxicity or cross-protection [18, 63]. In addition, our assumption of uniform relative biomass yields means that all communities in our model have a single stable equilibrium. How our results hold in the presence of alternative stable states and dynamical phases such as limit cycles is a question that remains to be answered [64, 65, 66, 67, 68, 69, 70]. Interestingly, we find that the assembly rule prediction also works well for trio communities with bistable pairs under three resources (SI Appendix V). Finally, we acknowledge that linear resource consumption is not the only possible mechanism for resource competition, and different models such as Monod model, Liebig’s law of the Minimum model, or a diauxic model may be more appropriate in some systems. Interestingly, we found that Monod model, Liebig’s law of the Minimum model, and the diauxic model all lead to community assembly that is often predicted by the same assembly rule [71](Fig. S4, SI Appendix I). In addition, different assumptions for resource competition such as fluctuating environment, community under oscillation or chaos instead of a single fixed point, or species directly modifying other competitors’ traits may lead to different bottom-up mappings between pairwise outcomes and community assembly. On the other hand, while we have focused on resource competition, there may be other ecological models that also lead to the experimentally observed bottom-up assembly rule. Future work is required to explore the extent of possible mechanistic origins behind the simplicity of community assembly.

Our approach highlights that, even when one uses mechanistic models, direct measurement of microscopic parameters is not always necessary for predictive power. This is a significant advantage of the empirical approach that has attracted many ecologists [72, 73, 67, 74, 75, 76, 30]. In the case of resource competition, fully characterizing the metabolism of each microbial species is a formidable experimental challenge due to many complications such as cross-feeding, diauxie, and co-utilization of resources. Our results illustrate how mechanistic insights can guide simple experimental predictions while circumventing the full microscopic complexity.

## Acknowledgements

The authors would like to thank the members of Gore Lab for comments on the manuscript. This work was supported by the Sloan Foundation and the Schmidt Polymath Award.

## Author contributions

HL, BB and JG conceived the study. HL performed theoretical modeling. HL and BB performed numerical simulations. All authors analyzed the data and wrote the manuscript.

## Methods

### Simulations of linear resource consumption model

We used simulations to complement our geometric approach with our resource competition model. In the main text, we used a linear resource consumption model described in Eq. 1 and Eq. 2. The simulations are implemented with Python 3.9.13.

For each condition (*S* species and *R* resources), we simulated 500 communities and numerically solved outcomes of all pairwise competitions and the all-species competition for each community. This was done by repeating simulations of 10 communities 50 times. For each simulation of 10 communities with *S* species and *R* resources (*S*ranging from 3 to 12, *R* ranging from 1 to 100), growth rates and supplied resource concentrations were drawn randomly in the following way.

Growth rates *r* _*μi*_ of each species *μ* on each resource *i* were sampled from an array of length 10*S* and approximately unit mean, 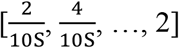 without replacement. The discretization ensured finite differences between growth rates of different species on a same resource and prevented numerical artifacts. Sampling from non-uniform, continuous distribution did not change the results (SI Appendix I.3). Note that growth rates on different resources were drawn independently, and there was no trade-off in growth rates. Results in the presence of trade-off are discussed in SI Appendix I.2.

Supplied resource concentrations 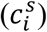 were drawn from uniform distribution between 0.1 and 1, and then were normalized to a fixed total concentration 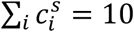. Biomass yields *Y*_*μi*_ were set to 1 for all species and resources.

Dilution rate was set to *δ* = 0.02. It is important to note that, due to the linearity of model, the only relevant scale of parameters is the scale between locations of ZNGIs and the supplied resource concentration, 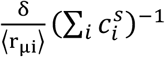 e.g. if all the ZNGIs and supplied resource concentration are scaled by the same factor, the relative position of supplied resource concentration to ZNGIs and consumption vectors would not change, and thus resource competition outcome would stay the same. Our choices of ⟨ rμ_i_⟩ = 1, δ = 0.02, and 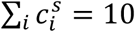 sets the supplied resource concentrations to be far from ZNGIs, which recapitulates laboratory microcosms and other resource-abundant environment in nature. We also found that different choice of δ = 0.2 did not change our results (SI Appendix I.4). This resource-abundant setup also implied that any extinction was driven by competition, not by dilution outpacing monoculture growth. With the same randomly generated array of growth rates and supplied resource concentrations, we simulated all 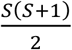 pairwise competitions and the all-species competition by running simulations with corresponding initial abundances (1 for species participating in competition, 0 for others). Finally, we implemented convex optimization to solve for the equilibrium of each competition [77, 78]. Species with population fraction < 10^5:^ were considered extinct.

### Quantification of agreement between simulation and prediction

In trio community simulations without cross-feeding, we compared the simulated community assembly with three different predictions: ‘Everyone survives’ that always predicts all three species to survive, ‘Fastest grower survives’ that predicts the best grower under the resource supply to exclude other two species, and the assembly rule. We quantified how well the simulated competitions agreed with each prediction by calculating the fraction of species in each trio that either survives while predicted to survive or goes extinct while predicted to go extinct. For example, if prediction tells that only A would survive and simulation results in only B surviving, the agreement is 0 for A, 0 for B, and 1 for C, resulting in an average 1/3 agreement for the trio.

In larger-community simulations and trio community simulations with cross-feeding, we compared the simulated community assembly to the assembly rule. The quantification followed the same protocol described above.

### Simulations of linear resource consumption model with cross-feeding

To implement simulations with cross-feeding, we considered the following equations:

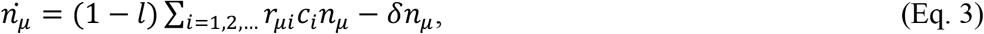

and

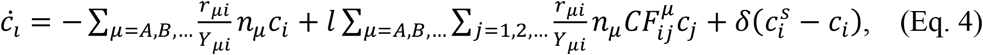

where *l* is the strength of cross-feeding, and 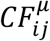 is the cross-feeding network between resources, which is nonzero if consumption of j-th resource produces *i*-th resource. For simplicity we set 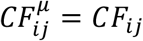 in our simulations. We assumed that each resource had probability *p* to cross-feed a less complex metabolite. Followingly, for each simulation of 10 communities, the cross-feeding matrix *CF*_*ij*_ was randomly generated as an upper triangle matrix:

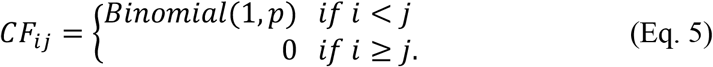

All other parameters were drawn in the same way as the simulations without cross-feeding. We implemented convex optimization to solve for the equilibrium of each competition.

## Data availability

Data files and simulation codes to reproduce all figures are available via Github (https://github.com/lachesis2520/resource_assembly_public).

## Footnotes

1 We note that such bottom-up prediction is not guaranteed even in purely pairwise models. For example, Lotka-Volterra model predict four trio outcomes under the same pairwise outcomes of two coexistences and one exclusion (Fig. 3C) [57].

2 Equivalently, the rule assumes that a species invades a community if and only if it can invade every member of the community. Yet another definition is that, a species collection coexists if all its sub-collections coexist [76].

3 This notion of hierarchy between coexistence across multiple scales has been recently formalized for any generic community assembly by Angulo et al.; in the formalism, this hierarchy is equivalent to the absence of disassembly holes [76].

