## Supplemental Information for "Resource competition can explain simplicity in microbial community assembly"

**This file includes:**

Figs. S1 to S15
Table S1
Appendix I
Appendix II
Appendix III
Appendix IV
Appendix V

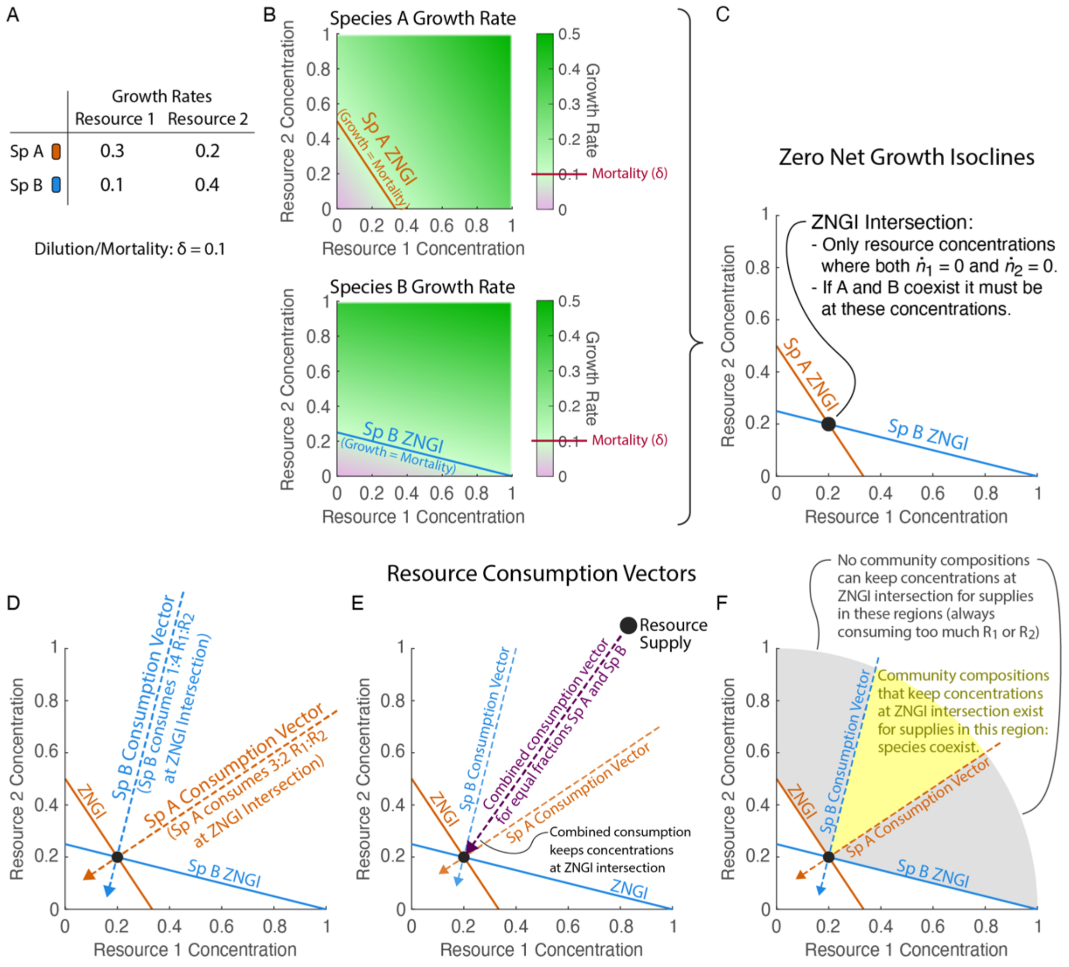


***Fig. S1. Additional explanation of the significance of ZNGIs and consumption vectors in the graphical approach.*** *(**A**) Growth rates for two species growing on two resources and the environmental dilution (i.e. species mortality) rate used as an example throughout this figure. (**B**) Species' growth rates as a function of resource concentrations shown using a colormap with the horizontal and vertical axes corresponding to the resource 1 and 2 concentrations. Species' Zero Net Growth Isoclines (ZNGIs, shown as lines on top of the colormap) are the resource concentrations at which a specific species' growth exactly matches the mortality rate. If a species survives with a constant population size, the steady-state resource concentration must lie on its ZNGI or else its population size would be overall growing or shrinking. (**C**) Species A and B's ZNGIs shown on the same plot with the horizontal and vertical axes again corresponding to the resource 1 and 2 concentrations. The ZNGIs intersect at a unique point, which is the only steady-state resource concentration at which both species will maintain constant population sizes. Therefore, if A and B coexist the steady-state resource concentration must be at the ZNGI intersection. (**D**) Plot from **C** with resource consumption vectors added as dashed lines. The slope of a consumption vector represents the ratio of how much resource 1 vs resource 2 each species is consuming at the ZNGI intersection. (**E**) Plot from **D** with an example resource supply drawn and the combined consumption vector of the community of species A and B that assembles under this resource supply shown as a purple dashed line. The consumption vector of the combined community always lies in between the consumption vectors of the individual species. At steady-state the combined consumption vector must point from the resource supply to the steady-state resource concentrations (which is necessary for the dilution and consumption to together exactly match the resource supply rate and for resource concentrations to therefore be constant). (**F**) Plot from **D** with the region of resource supplies that support the coexistence of species A and B highlighted in yellow and the regions of resource supplies that support the survival of either species in monoculture but not their coexistence shaded in gray. If the resource supply does not lie in between the species' individual consumption vectors, no community composition will have the necessary combined consumption to keep the resource concentrations at the ZNGI intersection and coexistence will not be possible.*

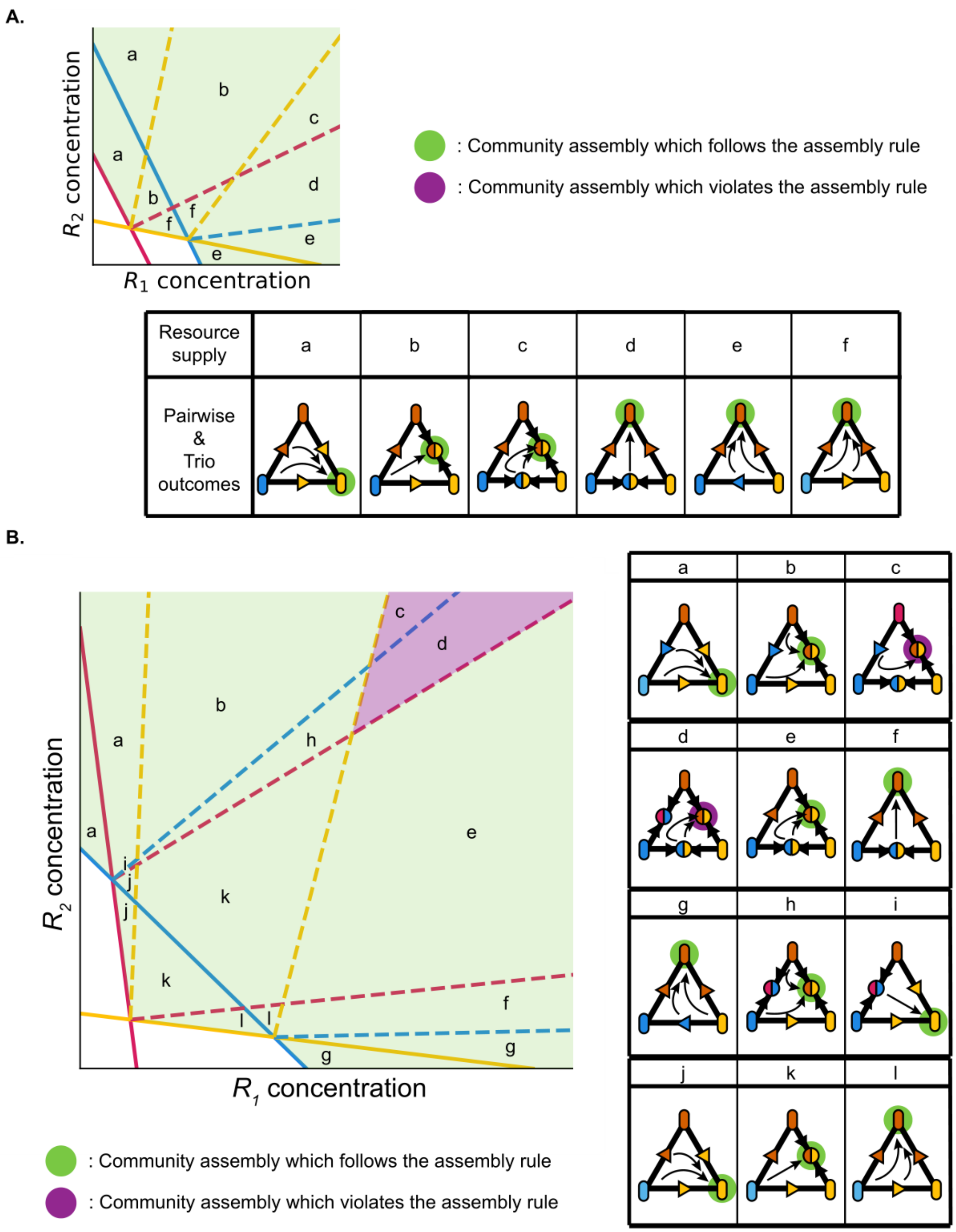


***Fig. S2. Community assembly of trios in Fig. 2 agrees with the assembly rule under most of resource supply concentration.*** *A. All 6 outcomes for the trio in Fig. 2A. Note that outcomes change only when resource supply moves over a resource consumption vector. All the outcomes follow the assembly rule. B. All 12 outcomes for the trio in Fig. 2B. Outcomes violate the assembly rule when resource supply is inside region c or d.*

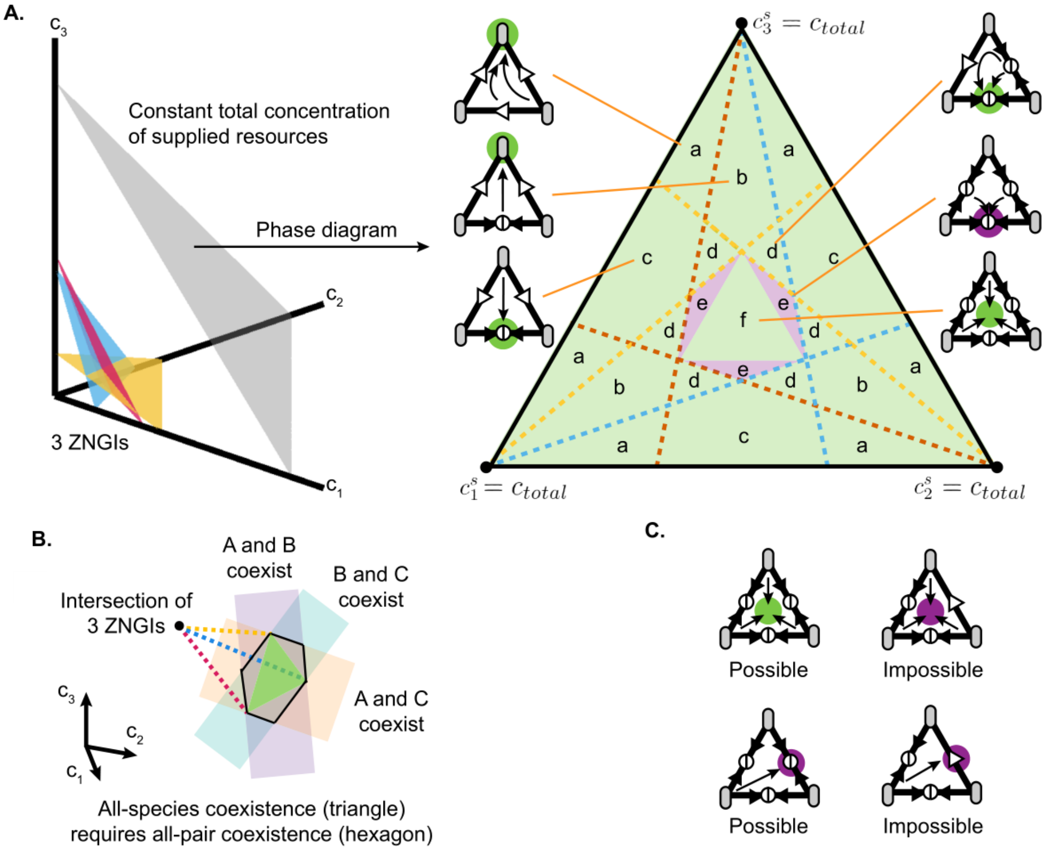


***Fig. S3. All-species coexistence requires all-pair coexistence, but not vice versa. A.*** *Geometric analysis of trio community competing for three resources illustrates all-pair coexistence both with and without all-species coexistence. In the 3-dimensional space of three resource concentrations, it is often useful to consider a simplex on which the total concentration of supplied resources is kept constant. The phase diagram of outcomes in the case of three ZNGIs intersecting at a point, projected on the simplex, is qualitatively same as the example illustrated on the right. We can see that the region where three species coexist in trio competition (region f) is inside the region where all pairs coexist (region e & f).* ***B.*** *Generically, the resource supply supporting all-species coexistence always supports all-pair coexistence as well, but not vice versa. This is because the convex hull of three resource consumption vectors, which is the region of all-species coexistence, is always included in the intersection of every region for each pairwise coexistence.* ***C.*** *This geometric constraint dictates that when a pair does not coexist it can never coexist in community assembly. This implies that the two outcomes on the left are possible while the other two on the right are impossible.*

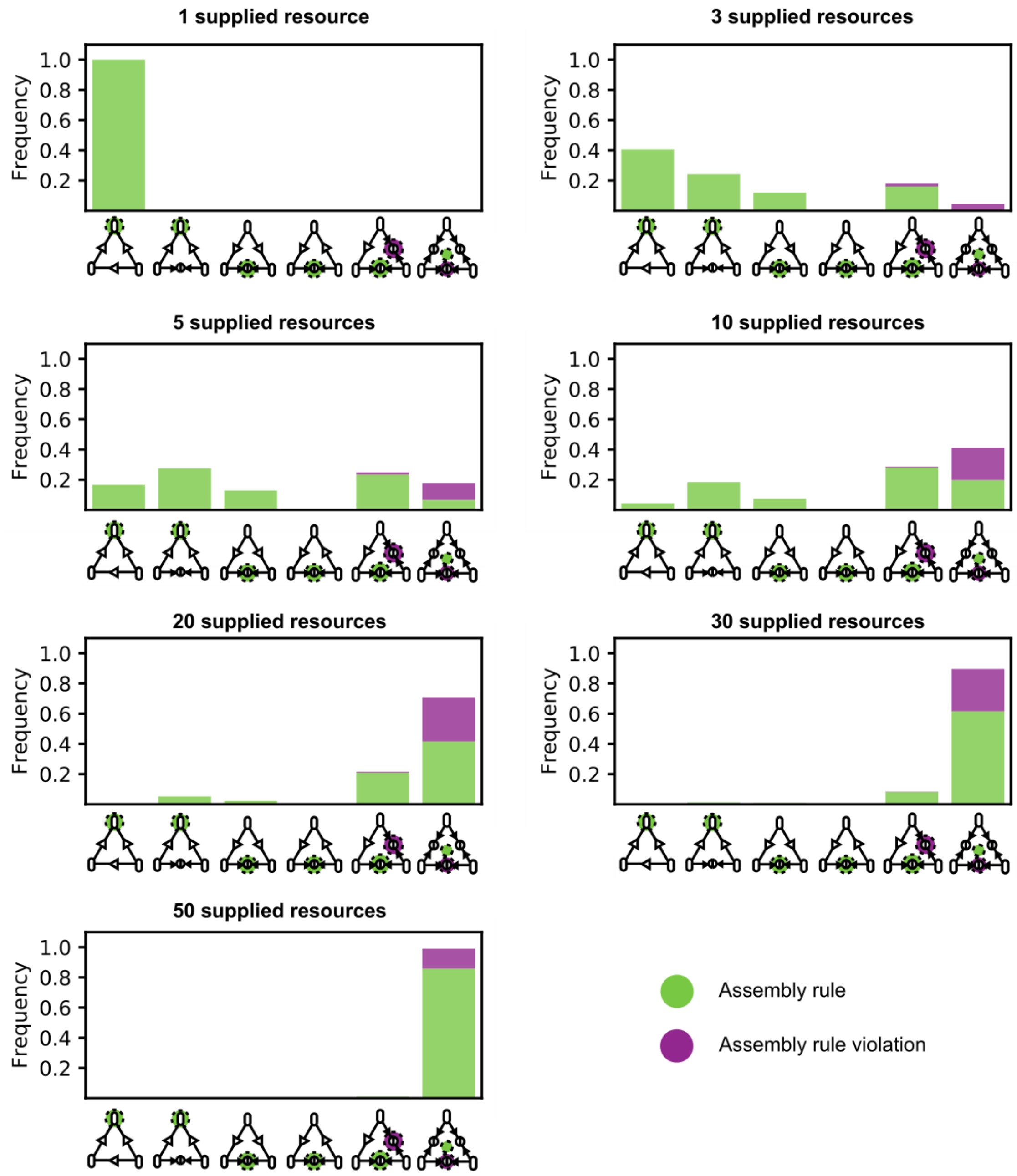


***Fig. S4. Trio community assembly outcomes for each number of supplied resources do not exhibit unexpected assembly rule violations.*** *In addition to 1, 5, and 50 resources cases shown in Fig. 4, here we show the distribution of outcomes under all number of resources that we have simulated. As the number of resources increase, species tend to coexist. Also, any violation observed in simulations is one of two violations that are expected from geometric analysis.*

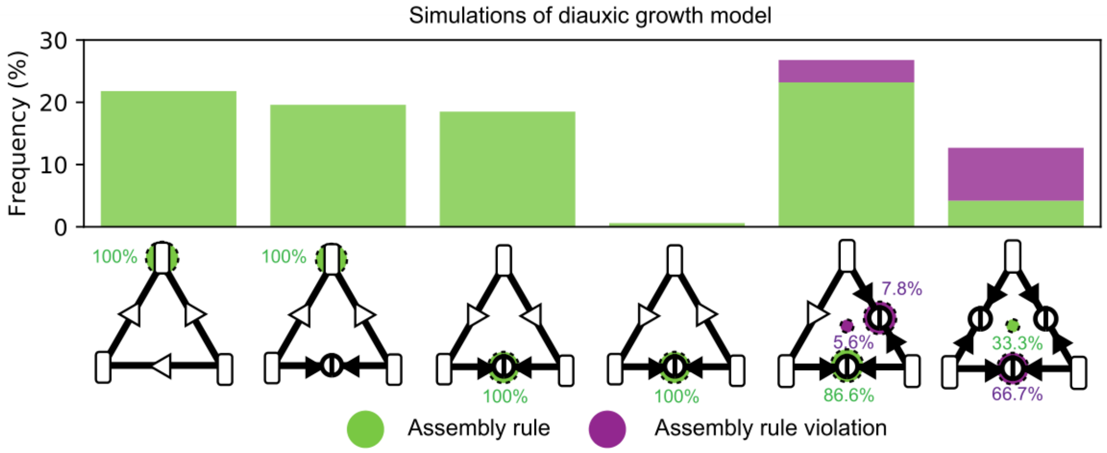


***Fig. S5. Trio community outcomes in diauxic growth model often agree with the assembly rule.*** *While our chemostat-like model in the main text assumes that species consume all resources simultaneously, microbes in nature often exhibit diauxic growth in which each resource consume one resource at a time, starting with its top preference and then moving on to its next remaining preference after each resource depletion. We tested how the assembly rule works under such growth dynamics by implementing a diauxic growth model used in Bloxham et al. [70]. Across 1000 simulated communities of three species growing on three resources, the assembly rule often predicted trio outcomes accurately from pairwise outcomes. (For each simulated community, each component of the resource supply was uniform-randomly sampled from 0 to 1 and the total resource supply was then normalized to 1, while growth rates were uniform-randomly sampled from 0 to 1. Each species' resource preference order matched the ordering of its growth rates such that each species first consumed whichever resource it grew fastest on. A dilution factor of 10 was used. Diauxic lags were assumed to be zero so that species instantly starts consuming the next preferred resource upon depletion of the previous resource.)*

| Number of resources | Communities with a non-intersecting ZNGI pair | Assembly rule prediction accuracy (%) | Communities without non-intersecting ZNGI pair | Assembly rule prediction accuracy (%) |
|---|---|---|---|---|
| 1 | 500 | 100 | 0 | - |
| 3 | 278 | 100 | 222 | 94.29 |
| 5 | 82 | 100 | 418 | 94.90 |
| 10 | 3 | 100 | 497 | 92.62 |
| 20 | 0 | - | 500 | 90.13 |
| 30 | 0 | - | 500 | 90.6 |
| 50 | 0 | - | 500 | 95.6 |

***Table S1. Assembly rule prediction always works perfectly for trio communities with non-intersecting ZNGI pairs.*** *From geometric analysis, we stated that the assembly rule should perfectly predict trio community assembly when any pair of ZNGI does not intersect. By inspecting the growth rates in each simulated community, we divide the dataset used in Fig. 4 into communities with and without non-intersecting ZNGI pairs. Here we show the number of communities in the two subsets and average assembly rule accuracy in each of the subsets. The assembly rule indeed perfectly works for simulated communities with non-intersecting ZNGI pairs, verifying our statement from geometric analysis.*

### Appendix I. Alternative models and parameters

#### I.1. Alternative models: summed Monod, Liebig's Law of the Minimum

In the main text, we considered a simple model of resource competition in which growth of a species is a sum of growth on each resource that is proportional to resource concentration. It is important to distinguish the results that hold under generic resource model framework from the results specific to the simple model that we considered.

The list of trio outcomes under two and three resources, identified by geometric analysis (Fig. 3B), is specific to the simple model we considered. While the same methodology of inspecting all possible phase diagrams can be applied to other resource competition models, the approach is limited to few numbers of resources and may not provide additional insights beyond the specific model. Given this limitation, we performed numerical simulations of two additional models to generalize results from the original simple model.

The first model is summed Monod model, which assumes type-II functional response between resource concentration and growth instead of linear relationship:

$$\dot{n}_\mu = \sum_{i=1,2,\ldots} r_{\mu i} \frac{c_i}{K_{\mu i}+c_i} n_\mu - \delta n_\mu, \qquad \textit{(Eq. S1)}$$

and

$$\dot{c}_i = -\sum_{\mu=A,B,\ldots} \frac{r_{\mu i}}{Y_{\mu i}} n_\mu \frac{c_i}{K_{\mu i}+c_i} + \delta(c_i^s - c_i), \qquad \textit{(Eq. S2)}$$

The second model is Liebig's Law of the Minimum model, which assumes that resources are never substitutable by one another:

$$\dot{n}_\mu = \min_{i=1,2,\ldots} r_{\mu i} \frac{c_i}{K_{\mu i}+c_i} n_\mu - \delta n_\mu, \qquad \textit{(Eq. S3)}$$

and

$$\dot{c}_i = -\frac{1}{Y_{\mu i}} \min_{i=1,2,\ldots} r_{\mu i} \frac{c_i}{K_{\mu i}+c_i} n_\mu + \delta(c_i^s - c_i), \qquad \textit{(Eq. S4)}$$

Next, we simulated these additional models in the following way.

For both models, as in the main text, we used the same sampling of growth rates $r_{\mu i}$ from uniform distribution between 0 and 2. This assumed that the maximal growth rates are orders of magnitude larger than the dilution rate $\delta = 0.02$. For both models, we also sampled Monod constants $K_{\mu i}$ from gaussian distribution with mean of 1 and standard deviation of 0.2. The summed Monod model is solved with convex optimization, and Liebig's Law of the Minimum model is solved with ODE solver.

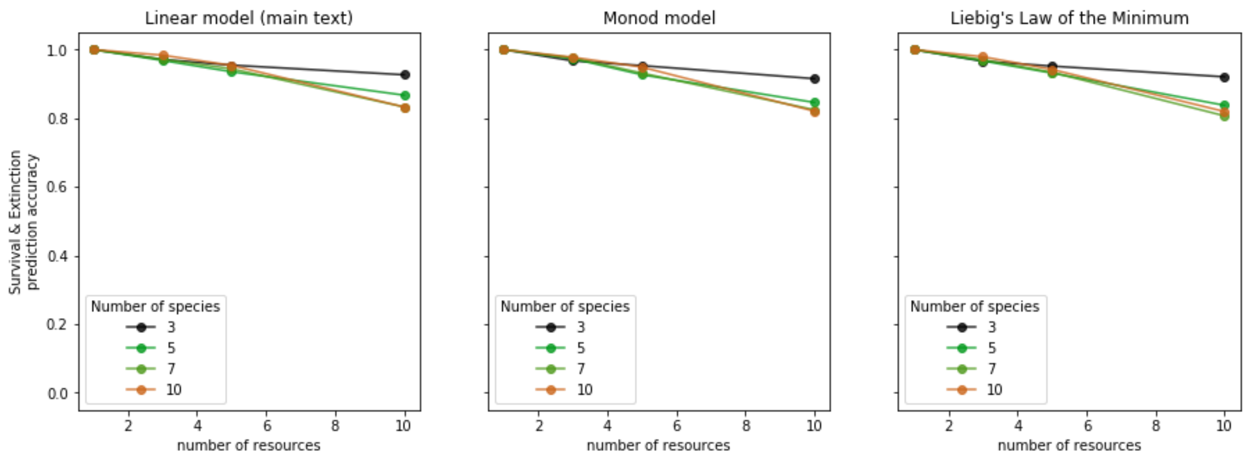


***Fig. S6. Alternative resource competition models often agree with the assembly rule.*** *Using the summed Monod model and Liebig's law of minimum model, instead of the linear model in the main text, did not significantly decrease the assembly rule prediction accuracy.*

We find that both the additional models also often lead to the simple community assembly as the assembly rule predicts.

#### I.2. Alternative parameters: metabolic tradeoffs

In the main text, we used uniformly distributed growth rates in simulations. While assembly rule violations in the simulations were rare, the frequency can depend on the underlying distribution of parameters. A particularly interesting scenario is metabolic tradeoff, which prevents a 'superorganism' that grows fast in all resources. Such metabolic tradeoff would increase the chance of ZNGI pairs crossing, which is a necessary condition for assembly rule violation (Fig. 2, Table S1).
To address this possibility further, we ran simulations with three different conditions:

1. No metabolic tradeoff (as in the main text)
2. Linear metabolic tradeoff $\sum_i r_{\mu i} = 1$
3. Quadratic metabolic tradeoff $\sum_i r_{\mu i}^2 = 1$

Interestingly, the second option (linear tradeoff) enables unlimited coexistence that violates our claim under a generic resource competition that all-species coexistence requires coexistence of all pairs (Fig. S3). While this scenario requires fine-tuned parameters, it can then lead to frequent violations of the assembly rule.

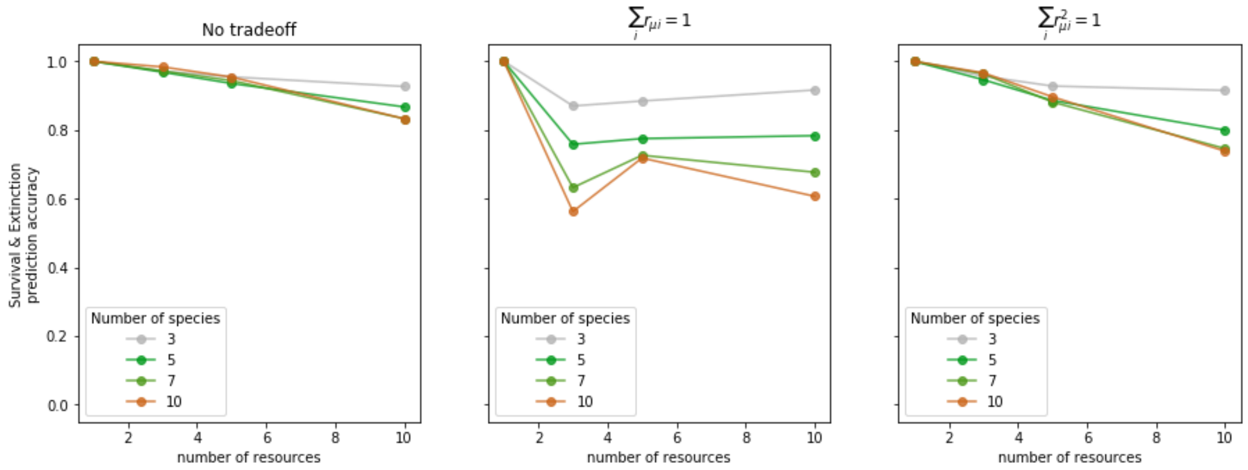


***Fig. S7. Resource competition under metabolic tradeoff often follow the assembly rule unless fine-tuned.*** *A fine-tuned linear tradeoff dropped the assembly rule prediction accuracy, while a quadratic tradeoff did not significantly decrease the assembly rule prediction accuracy.*

We can see that, as expected, a linear tradeoff leads to a frequent violation of the assembly rule. Nevertheless, the quadratic tradeoff does not significantly increase the frequency of assembly rule violations, suggesting that the large drop of accuracy under the linear tradeoff is a fine-tuned special case. In the end, the frequency of assembly rule violation increases as more ZNGI pairs cross, and this may be associated with evolution, history of coexistence, and metabolic tradeoffs.

#### I.3. Alternative parameters: gaussian and exponential distributions of growth rates

After studying the effect of metabolic tradeoff, we also considered different choices of parameter distribution.

First, instead of assuming that all species can consume all resources, we simulated 'specialist' scenario in which each species consume only one random resource. We also simulated an intermediate scenario in which each species has 50% chance to consume each resource.

We also ran additional simulations with growth rates that are not uniformly distributed. We simulated gaussian distribution (( $p(r_{\mu i}) \sim e^{-\frac{(1-r_{\mu i})^2}{2\times 0.2^2}}$ ) and exponential distribution ($p(r_{\mu i}) \sim e^{-r_{\mu i}}$) with the same mean growth rate 1 as the uniform distribution used in the main text.

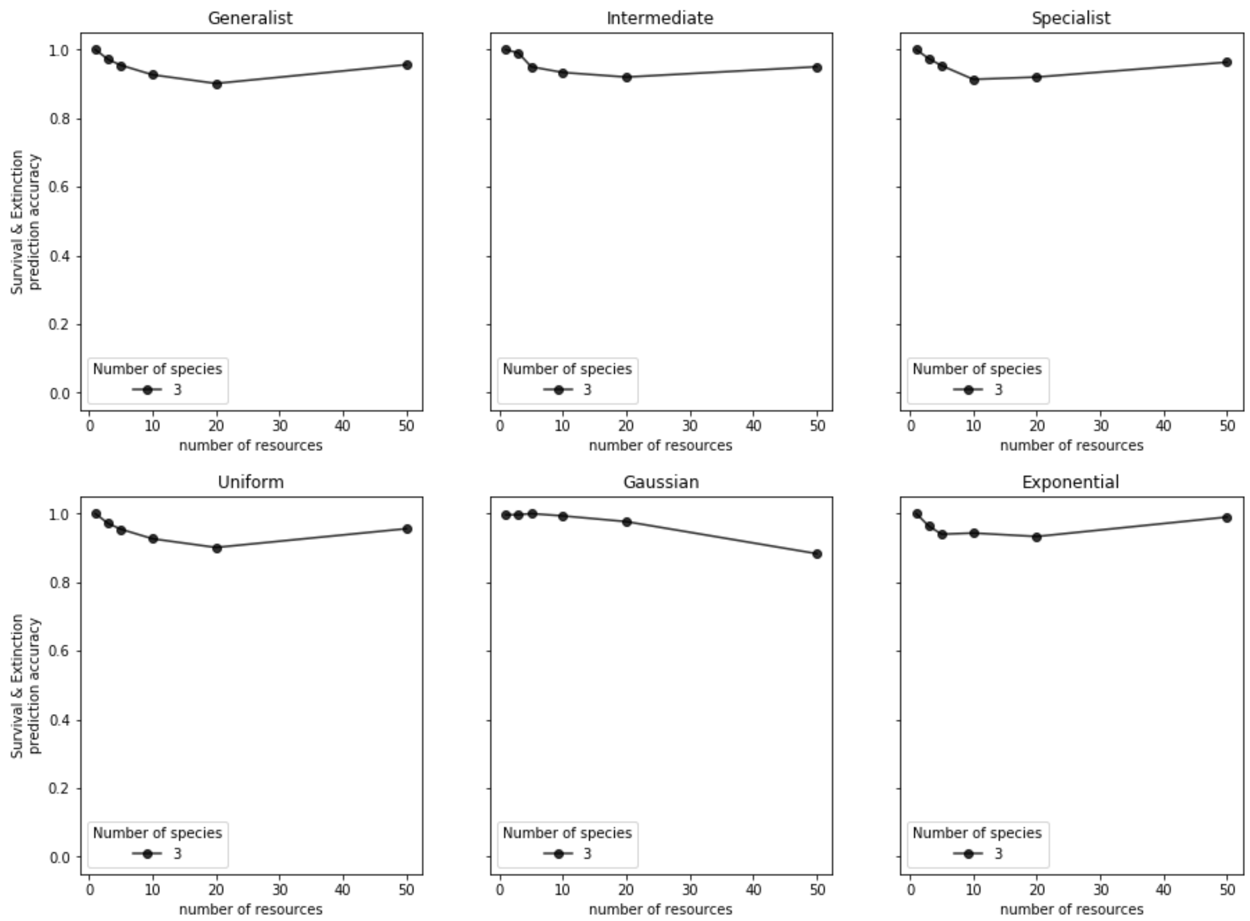


***Fig. S8. Alternative choices of growth rates often agree with the assembly rule.*** *The assembly rule prediction accuracy did not decrease significantly when species did not grow on all resources and when growth rates were not uniformly distributed.*

In all these simulations, we did not find significant decrease in the accuracy of assembly rule predictions.

### I.4. Alternative parameters: smaller scale of growth rates compared to dilution rates

In the simple model we considered in the main text, locations of ZNGIs are controlled by the ratio between growth rates and dilution rates, i.e. $\delta / r_{\mu i}$ . Thus, both decrease growth rates or increase in dilution rate equivalently realize environment with less supply resource concentration.

To test if this scale between growth and death may change the assembly rule prediction accuracy, we ran additional simulation with 10-fold increase in dilution rates.

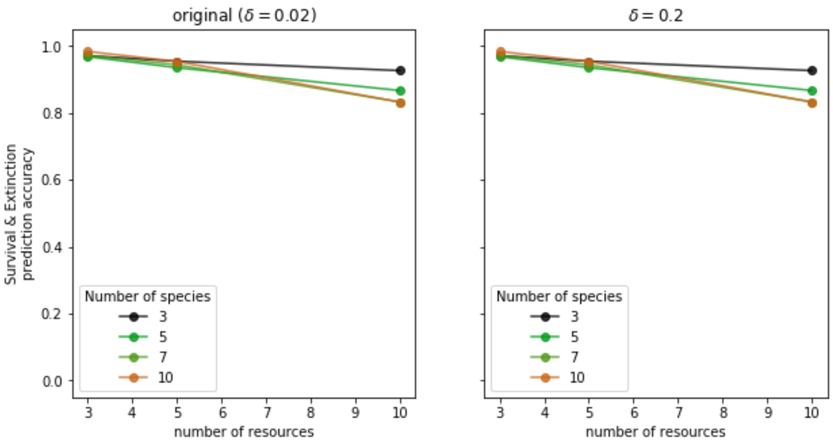


***Fig. S9. Resource competition under stronger disturbance often agree with the assembly rule.*** *The assembly rule prediction accuracy did not decrease significantly when dilution rate was increased by 10-fold.*

The simulations show that the assembly rule prediction works well even when dilution rate is less than an order of magnitude apart from growth rates. Even larger dilution rate may lead to frequent washout of all species in competitions, in which case assembly rule may not be well-defined. We do not focus on this scarce resource regime, as a broad range of natural environment and most of laboratory experiment have much larger supplied resource concentration than the minimum concentration required for single species survival.

### I.5. Alternative parameters: cross-feeding with varying numbers of species and resources

While we focused on exploring all regimes of cross-feeding from sparse to ubiquitous and from weak to strong in the main text, we missed out another axes of complexity, especially the number of species. To address this, here we show how assembly rule predictions work for simulations with different sized assemblages (3, 5, 7, and 10 species) and different number of cross-fed metabolites (2, 4, and 9 metabolites and a single supplied resource). Both the leakage rate and probability for a resource to release each other resource are set to 0.5. In addition, while our main text simulation assumed hierarchical resources and one-way cross-feeding from upstream to downstream, we also simulated two-way cross-feeding from each resource to any other resources. The two-way cross-feeding is implemented by sampling not only the upper triangle but also the lower triangle elements of cross-feeding matrix from binomial distribution. Specifically, the cross-feeding matrix was generated as:

$$CF_{ij} = \begin{cases} Binomial(1,p)/R & if\ i \neq j \\ 0 & if\ i = j. \end{cases} \qquad \text{(Eq. 5)}$$

The additional factor of $1/R$ ensured that cross-feeding does not break energy conservation. This was not required in one-way cross-feeding since biomass yields could be arbitrarily rescaled.

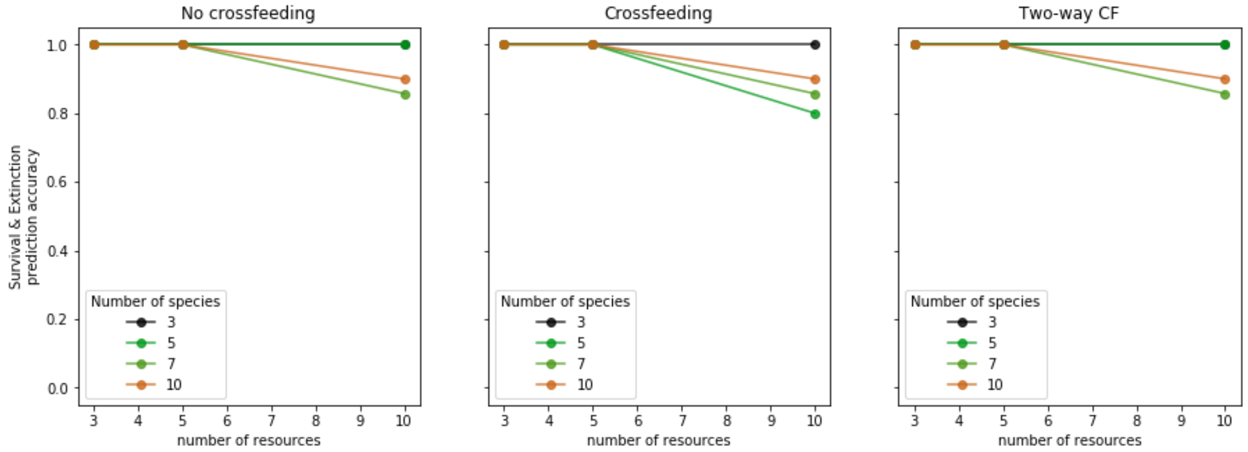


***Fig. S10. Resource competitions with cross-feeding often agree with the assembly rule.*** *Across varying number of species and resources, the assembly rule prediction accuracy did not decrease significantly when either hierarchal or fully connected cross-feeding networks were present.*

The simulations show that the existence of cross-feeding does not significantly affect the assembly rule prediction accuracy over varying number of species and resources.

### Appendix II. Trio community assembly under two supplied resources

In this appendix, we find all cases of 3-species community assembly under two resource supply. Assuming biomass yields are uniform or have the structure $Y_{\mu i} = a_\mu y_i$ (as in the Main Text), all possible scenarios for trio community can be categorized by how their ZNGIs intersect. There are 6 different ways for three lines (each with positive intercepts on both axes) to intersect on the quarter-plane of two positive resource concentrations. And for each topology of ZNGIs, we obtain a phase diagram of how different region of supplied resource concentrations lead to different pairwise and trio outcomes.

We also note that there are some important constraints on the resource consumption vectors:

First, two consumption vectors of a species from different points of its ZNGI can never cross. To see this, let us consider any two points $(\mathrm{c}_1^\mathrm{a}, \mathrm{c}_2^\mathrm{a})$ and $(\mathrm{c}_1^\mathrm{b}, \mathrm{c}_2^\mathrm{b})$ on a single ZNGI, Then consumption vectors from them have slopes $\frac{Y_{\mu 1} r_{\mu 2}}{Y_{\mu 2} r_{\mu 1}} \frac{\mathrm{c}_2^\mathrm{a}}{\mathrm{c}_1^\mathrm{a}}$ and $\frac{Y_{\mu 1} r_{\mu 2}}{Y_{\mu 2} r_{\mu 1}} \frac{\mathrm{c}_2^\mathrm{b}}{\mathrm{c}_1^\mathrm{b}}$, respectively. The slope will always be steeper for the point with larger $\frac{\mathrm{c}_2}{\mathrm{c}_1}$, and since the two points lie on the same line with a negative slope, the two consumption vectors will never intersect.

Second, by assuming that the biomass yield $Y_{\mu i} = a_\mu y_i$, we find a similar relation between consumption vectors of different species. Following the same analysis, we can see that the slopes will be steeper for larger $\frac{r_{\mu 2}}{r_{\mu 1}} \frac{\mathrm{c}_2}{\mathrm{c}_1}$. Thus by inspecting the slopes of ZNGIs and the points of intersections, we can tell whether two consumption vectors may or may not intersect.

For the sake of clarity, here we show the phase diagrams for all 6 topologies. For each arrangement of ZNGIs, a phase diagram with the maximum number of outcomes is shown.[1]

[1] By adjusting the yields and exact locations of the ZNGI intersections it is possible to not have all possible regions for each ZNGI arrangement. The yields and exact intersections in these plots where chosen to have all possible regions occur. (The constrain $Y_{\mu i} = a_\mu y_i$ prevents consumption vectors from being sufficiently angled to create additional regions that the reader might otherwise imagine.)

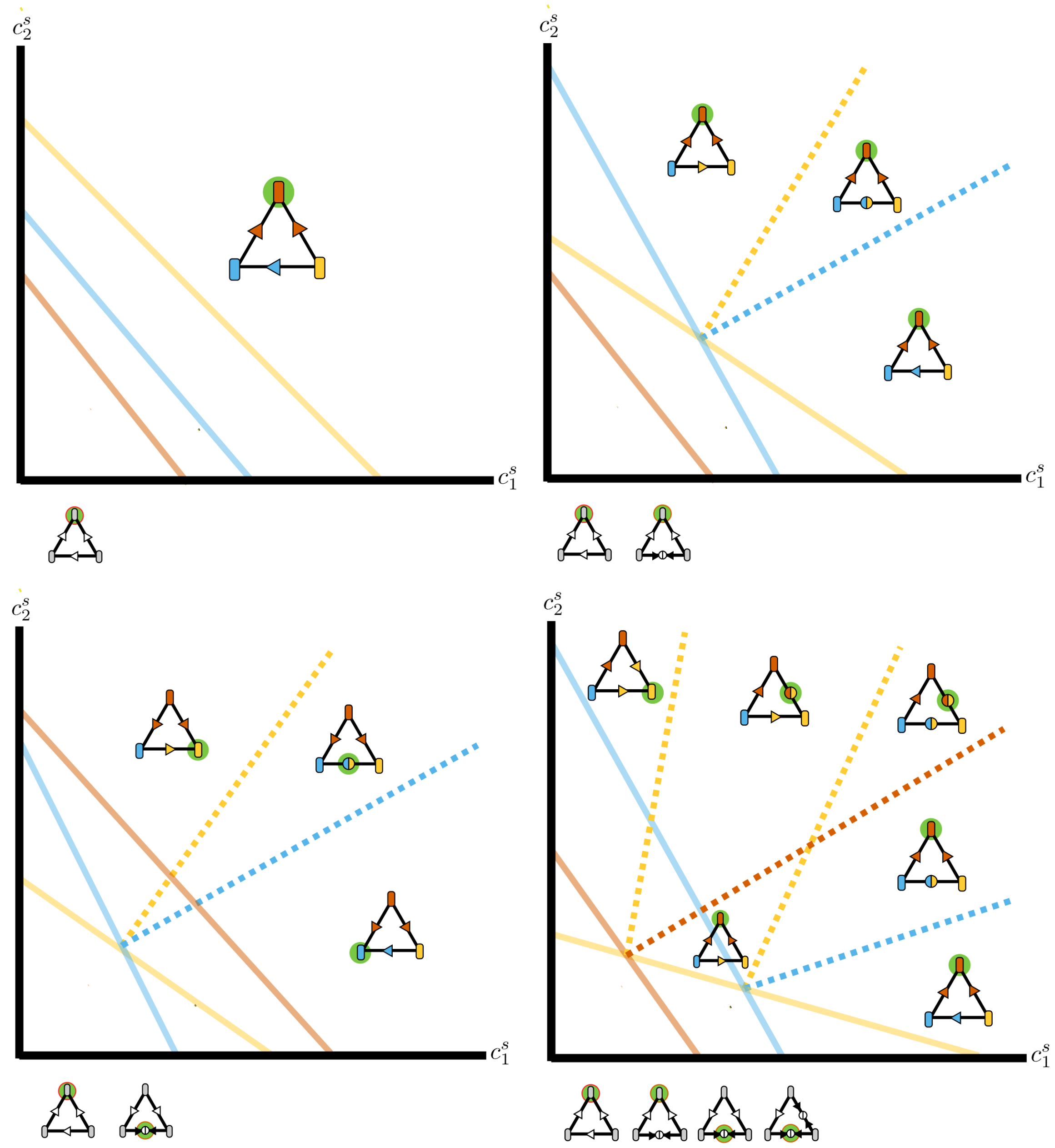
$c_2^s$
$c_1^s$

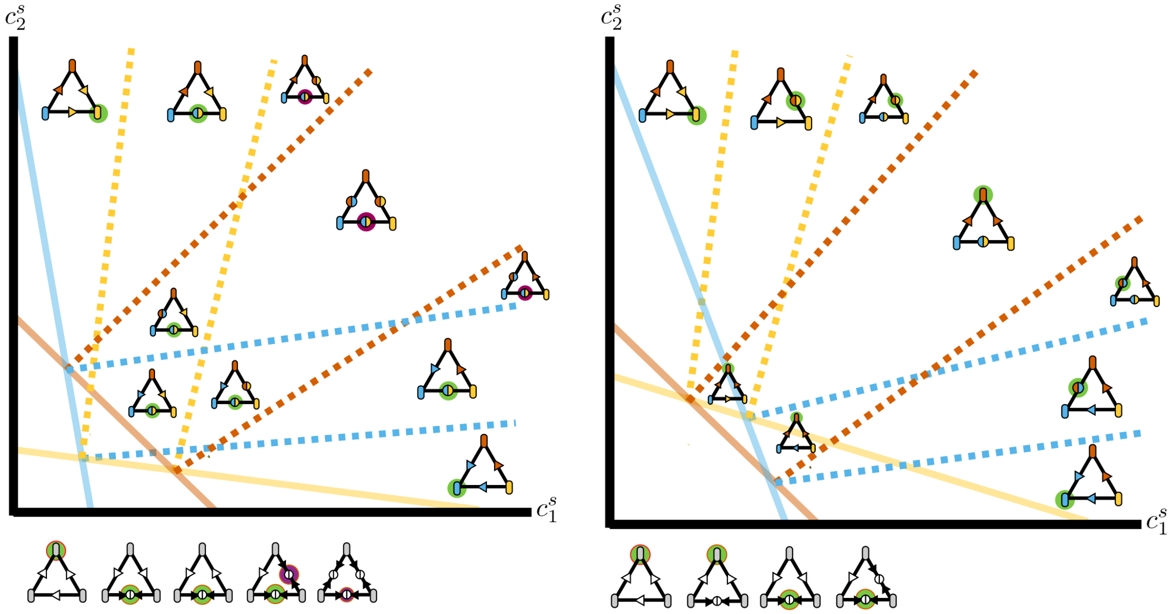


Fig. S11. Phase diagrams for all ZNGI topologies of 3 species on 2 resources. The bottom left diagram can lead to two distinct violations (one of which appears twice due to symmetry). All other cases always follow the assembly rule.

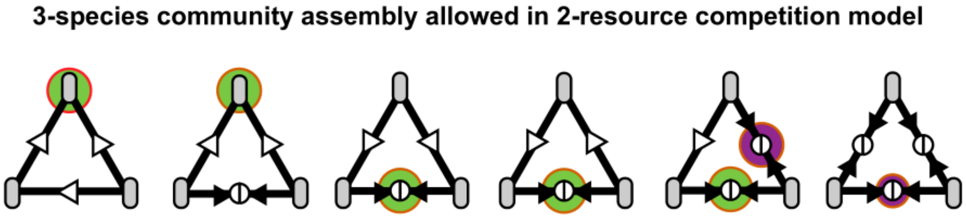


Fig. S12. The complete list of 3-species community assembly under two resources. Compared to the three-resources case (Fig. 4B), this list does not include three-species coexistence under three pairwise coexistence.

### Appendix III. Trio community assembly under three supplied resources

In this appendix, we find all cases of 3-species community assembly under three resource supply. Similar to the previous case, we identify all possible ZNGI arrangements and corresponding phase diagrams. As each ZNGI is a plane in this three-dimensional case, two ZNGIs intersect along a line and three ZNGIs at a single point may intersect. By counting such intersections that either lie on the innermost ZNGIs or lie outside them, we find all 7 possible ZNGI arrangements and possible community assembly under each of them (While we do not present a mathematically rigorous proof but instead focus on providing a more intuitive understanding, we are unaware of any other community assembly being possible).

Remarkably, many ZNGI arrangements in 3D are equivalent to 2D arrangements; one may find an axis along which all 2D slices of ZNGI planes have the same arrangements (such that rotating the 2D slice about the axis does not change the qualitative arrangement of the ZNGI projections on the slice). When this happens, as in the illustrated example on the left, since the 3D ZNGI arrangements is equivalent to a trivial extension of 2D arrangement along a new axis, the set of pairwise and trio outcomes is the same as its equivalent 2-resource competition's.

Given this, let us examine each of all 7 phase diagrams.

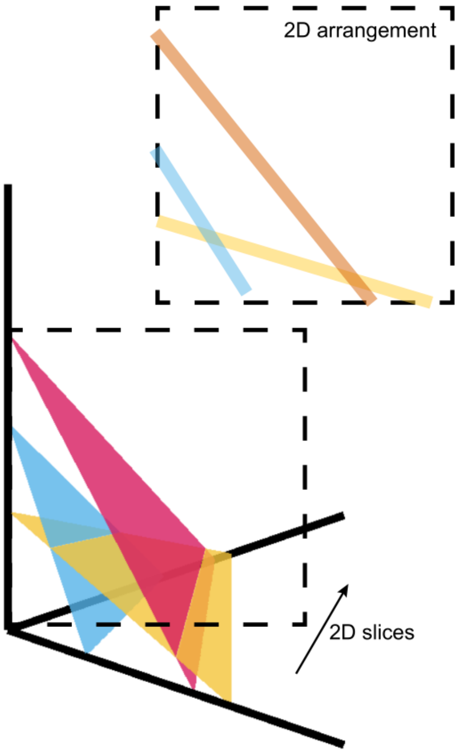

1. 0 ZNGI intersections

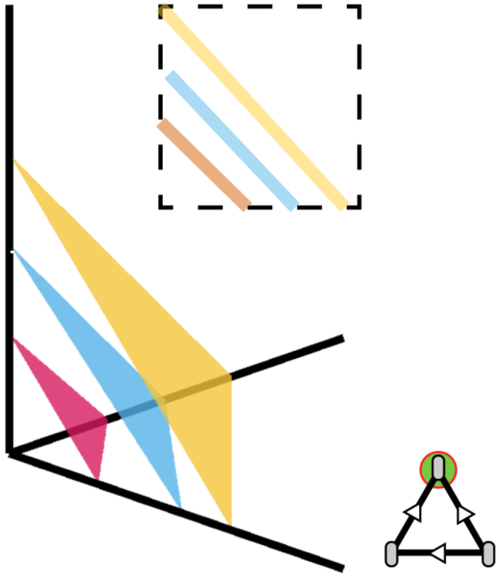

This case is equivalent to the 2-resource case with no ZNGI intersections. Fast grower excludes others in all competitions, and only one set of pairwise and trio outcome is possible.

2. 0 intersection on innermost ZNGIs, 1 intersection outside

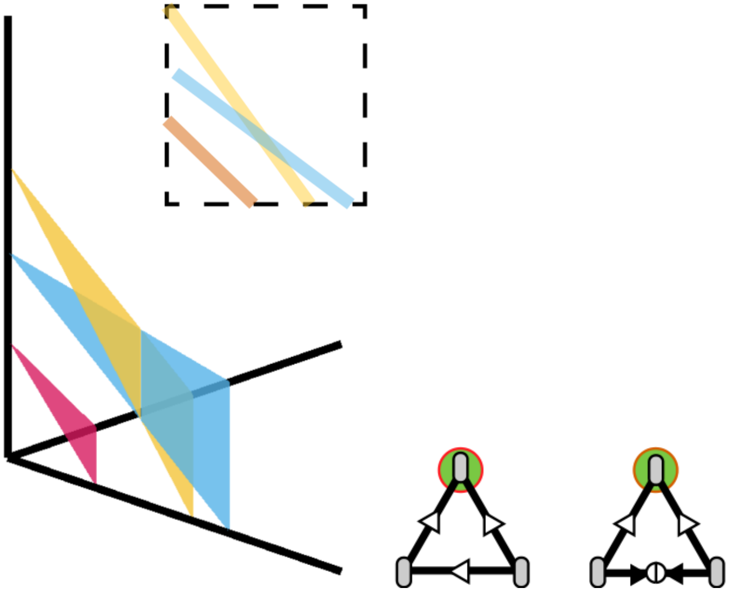

This case is equivalent to the 2-resource case with one ZNGI intersection outside the innermost ZNGI. The fastest grower (red) excludes others, and the others may coexist or exclude one another depending on resource supply.

3. 1 intersection on innermost ZNGIs, 0 intersection outside

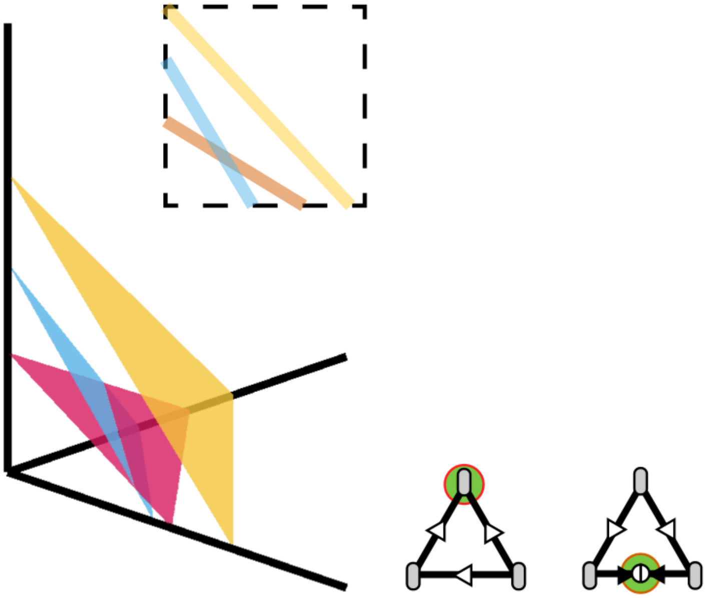

This case is equivalent to the 2-resource case with two innermost ZNGIs intersecting. The slowest grower (yellow) is excluded by others, and the others may coexist or exclude one another depending on resource supply.

4. 1 intersection on innermost ZNGIs, 1 intersection outside

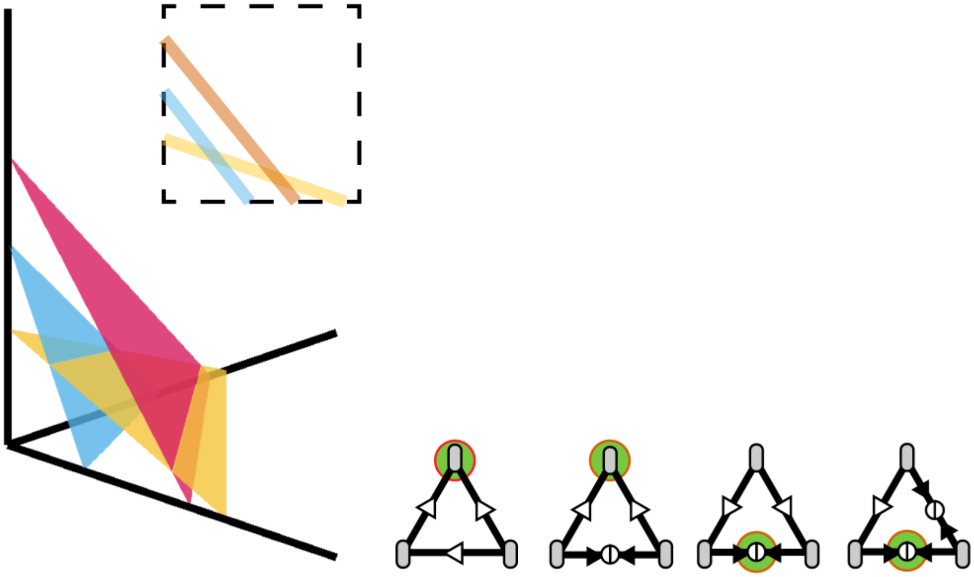

This case is equivalent to the 2-resource case with one ZNGI intersecting with the other two.

5. 1 intersection on innermost ZNGIs, 2 intersections outside

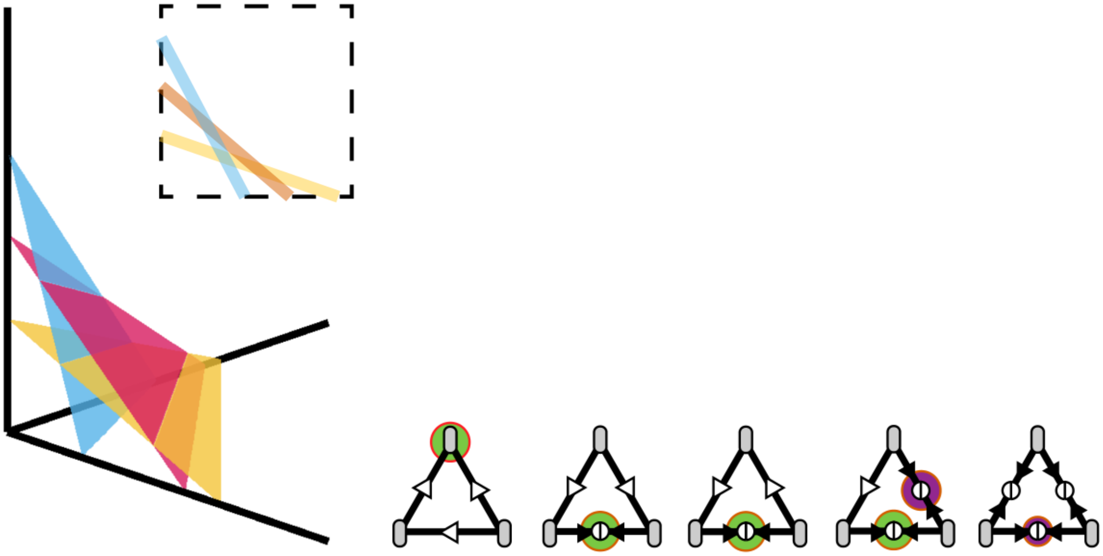

This case is equivalent to the 2-resource case with all ZNGI pairs intersecting, with innermost ZNGIs consisting of two ZNGIs.

6. 2 intersections on innermost ZNGIs, 1 intersection outside

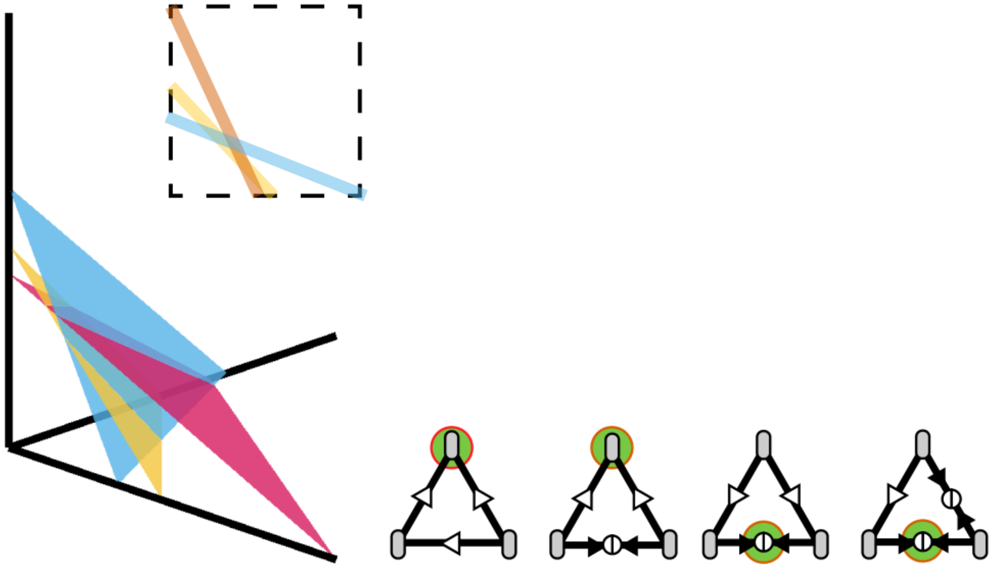

This case is equivalent to the 2-resource case with all ZNGI pairs intersecting, with innermost ZNGIs consisting of three ZNGIs.

7. Three ZNGIs intersect at a point

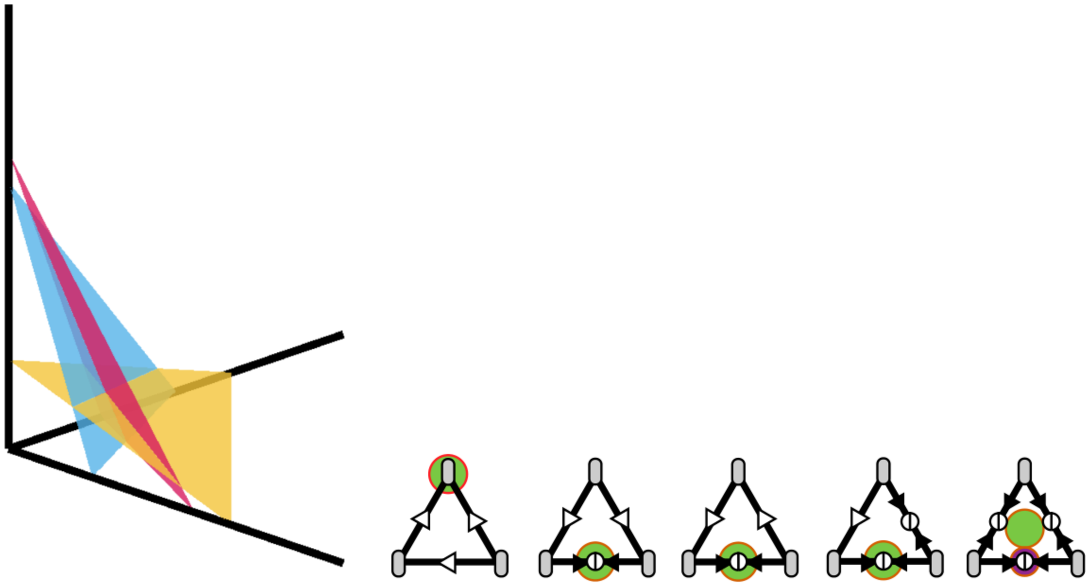

This case is not equivalent to any of 2-resource cases, which we can immediately see from the fact that three ZNGIs cannot intersect at a point on a plane in general. Remarkably, this is the only case where all three species can coexist. The phase diagram of outcomes for this arrangement is illustrated in Fig. S2.

Overall, we expect 8 possible 3-species community assembly, as shown in Fig. 4B in the main text.

### Appendix IV. Harmonic rule as a quantitative assembly rule

In the main text, we proposed 'harmonic rule' as a quantitative version of the assembly rule, which predicts a species' fraction in community assembly as a harmonic mean of the species' fraction in its pairwise competitions between other surviving species.

In this supplement, we discuss the origin of this rule in more details.

Let us consider a scenario where any species consume only one resource, i.e. all species are resource specialists. Then only the fastest grower for each available resource survives. Moreover, when a species survives, its population size at equilibrium is independent of its competitor. Specifically, a species' abundance can be easily calculated as:

$$n_\mu = \begin{cases} Y_{\mu i}\left(c_i^S - \frac{\delta}{r_{\mu i}}\right) & if\ r_{\mu i} = \max_\nu r_{\nu i} \\ 0 & otherwise \end{cases} \qquad \text{(Eq. S1)}$$

While the specific form of $n_\mu$ depends on the choice of resource competition model, the following results generally hold as long as $n_\mu$ does not depend on its competitors. For example, both the summed Monod model and Liebig's Law of the Minimum model in Appendix I obey the following results as long all species are resource specialists.
First, let's assume that species A survives in community assembly. Then it should be the fastest grower for its resource, and thus it survives in all pairwise competitions. Also, any surviving species in community assembly should consume another resource, and thus it should coexist with species A in a pairwise competition too. With this in mind, we can rewrite the fraction of species A in community assembly as:

$$f_A = \frac{n_A}{\sum_\mu n_\mu} = \frac{1}{\sum_\mu \frac{n_\mu^{A\mu}}{n_A^{A\mu}}} = \frac{1}{1+\sum_{\mu\neq A}\left(\frac{1}{f_A^{A\mu}}-1\right)}, \qquad \text{(Eq. S2)}$$

where $f_A$ is the fraction of species A in community assembly, index $\mu$ runs over only the surviving species in community assembly, and $f_A^{A\mu}$ is the fraction if species A in pairwise competition between species A and $\mu$.

Next, let's assume that species A does not survive in community assembly. Then it should not be the fastest grower for its resource, and there exists another species B which grows fastest on the resource, survives in community assembly, and excludes species A in pairwise competition. This implies that $f_A^{AB} = 0$ and $f_A = 0$. Thus, Eq. S2 works for any species A regardless of its survival in community assembly.

Now we have a 'modified harmonic rule' which should work perfectly for specialist communities:

$$f_A = \frac{1}{1+\sum_{\mu\neq A}\left(\frac{1}{f_A^{A\mu}}-1\right)}. \qquad \text{(Eq. S3)}$$

This equation resembles harmonic mean $\left(\left\langle \frac{1}{f_A^{A\mu}} \right\rangle_{\mu\neq A}\right)^{-1}$. We can see write down the relationship between this modified harmonic rule and harmonic mean behaves as a function of $S$, the number of surviving species in community assembly.

$$\frac{f_A}{\left(\left\langle \frac{1}{f_A^{A\mu}} \right\rangle_{\mu\neq A}\right)^{-1}} = \frac{1}{S-1-(S-2)\left(\left\langle \frac{1}{f_A^{A\mu}} \right\rangle_{\mu\neq A}\right)^{-1}}. \qquad \text{(Eq. S4)}$$

We can see that the deviation between two estimations is zero when $S = 1$ and $S = 2$. As $S$ increases further, $f_A$ becomes smaller than $\left(\left\langle \frac{1}{f_A^{A\mu}} \right\rangle_{\mu\neq A}\right)^{-1}$ and $\lim_{S\to\infty} f_A = 0$.

To compare between the performance of two estimates, we applied them to simulated communities.

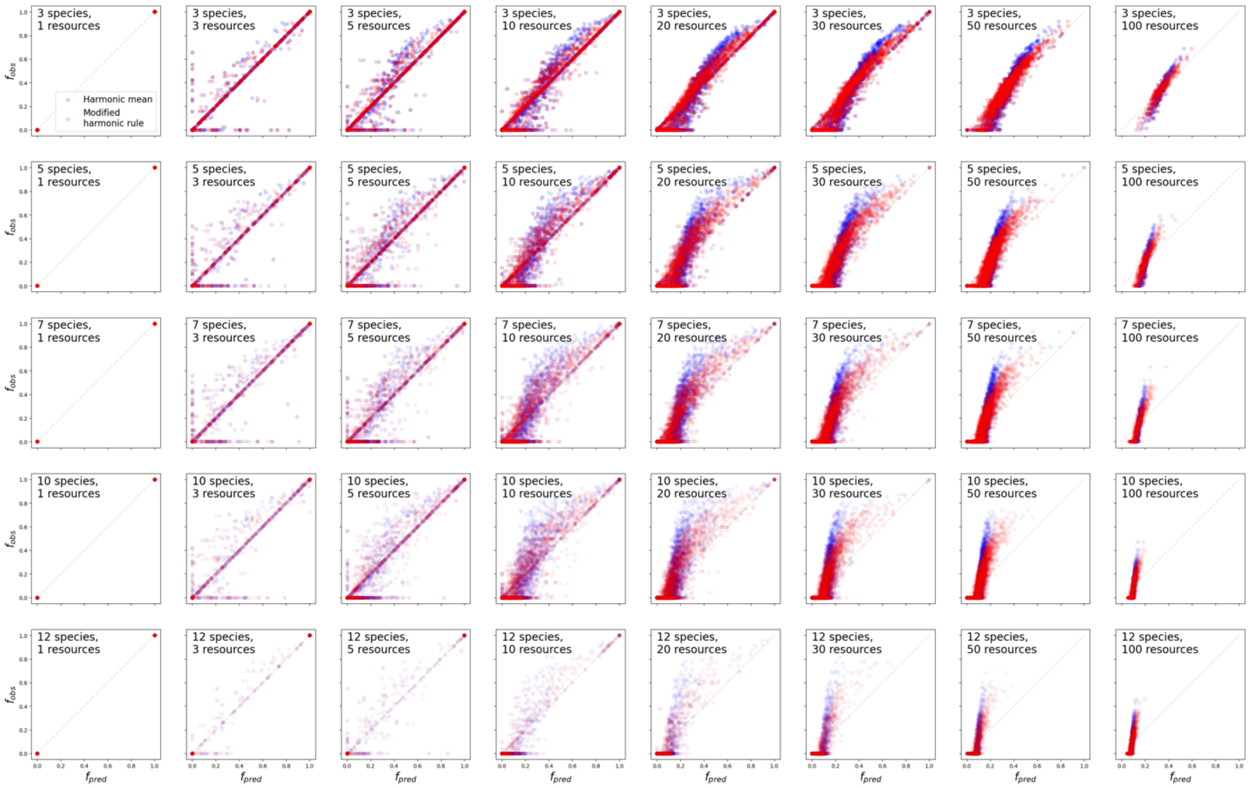


***Fig. S13. Modified harmonic rule works better than harmonic mean for predicting resource competition outcomes.*** *With many species and resources, the modified harmonic rule prediction results in smaller systematic deviation from observed species than harmonic mean prediction does.*

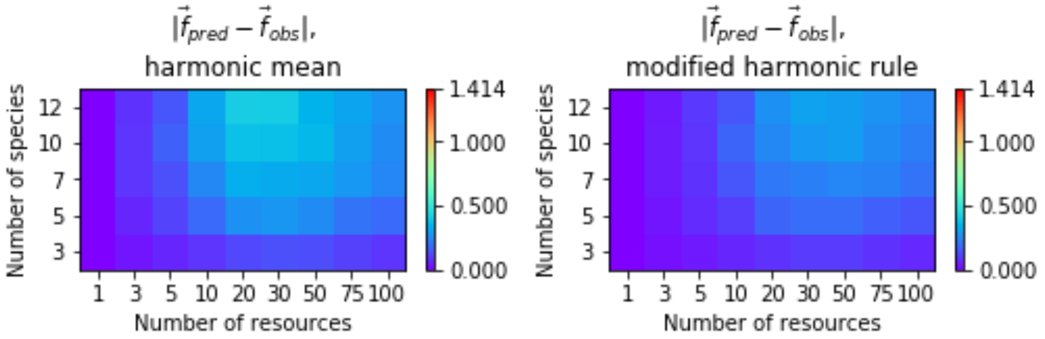


***Fig. S14. Modified harmonic rule works better than harmonic mean for predicting resource competition outcomes.*** *With many species and resources, the modified harmonic rule prediction results in smaller L2-distance from simulated community compositions than harmonic mean prediction does.*

We find that both modified harmonic rule and simple harmonic mean predictions work well for communities with few (5 or less) resources or 3 species. While trivial cases of one- or two-species survivals, in which cases both estimates should hold perfectly, may have been reflected in the trends, it is still interesting that such simple estimates can make bottom-up predictions fairly well. For more complex communities, both estimates start to underestimate fractions of surviving species and also predict false survivals. In this regime modified harmonic rule works better than the harmonic mean.

### Appendix V. Resource dynamics with bistable pairs recapitulate the assembly rule

In the main text, we assumed that all biomass yields are the same, independent of species and resources. As a result, each pair of species could either exclude one another or coexists and the pairwise outcomes were independent of initial species abundance. In general pairwise co-culture can also lead to bistability, in which the outcome depends on the initial species abundances. Each species cannot invade the other species and, as a result, initially abundant species (relative to the threshold set by pairwise interaction) excludes its competitor. This bistability is important for many ecological phenomena such as alternative stable states and priority effect, and thus warrants this additional investigation.

In this appendix, we extend our analysis by studying communities with one or more bistable pairs. We first explain how the assembly rule predicts community outcomes in the presence of bistable pairs. Next, we discuss the condition for pairwise bistability under our resource-consumer model. Finally, we demonstrate that simulations of our resource model with bistable pairs agree with the assembly rule predictions. This reinforces our main text results that resource dynamics can explain the assembly rule.

#### V.1. Assembly rule with bistable pairs

The assembly rule predicts that species A will exclude species B in community assembly if A excludes B in pairwise competition [34]. Equivalently, the rule assumes that a species invades a community if and only if it can invade every member of the community. This definition based on invasibility can be readily applied to communities with bistable pairs. When A and B are bistable, A and B cannot invade each other. On the other hand, when A and B coexists, A and B can invade each other. Finally, when A excludes B, then A can invade B while B cannot invade A.

We can apply this definition and extend assembly rule predictions to community assembly in the presence of bistable pairs. The complete list of assembly rule prediction for trio, an extension compared to Fig. 1A, is shown below.

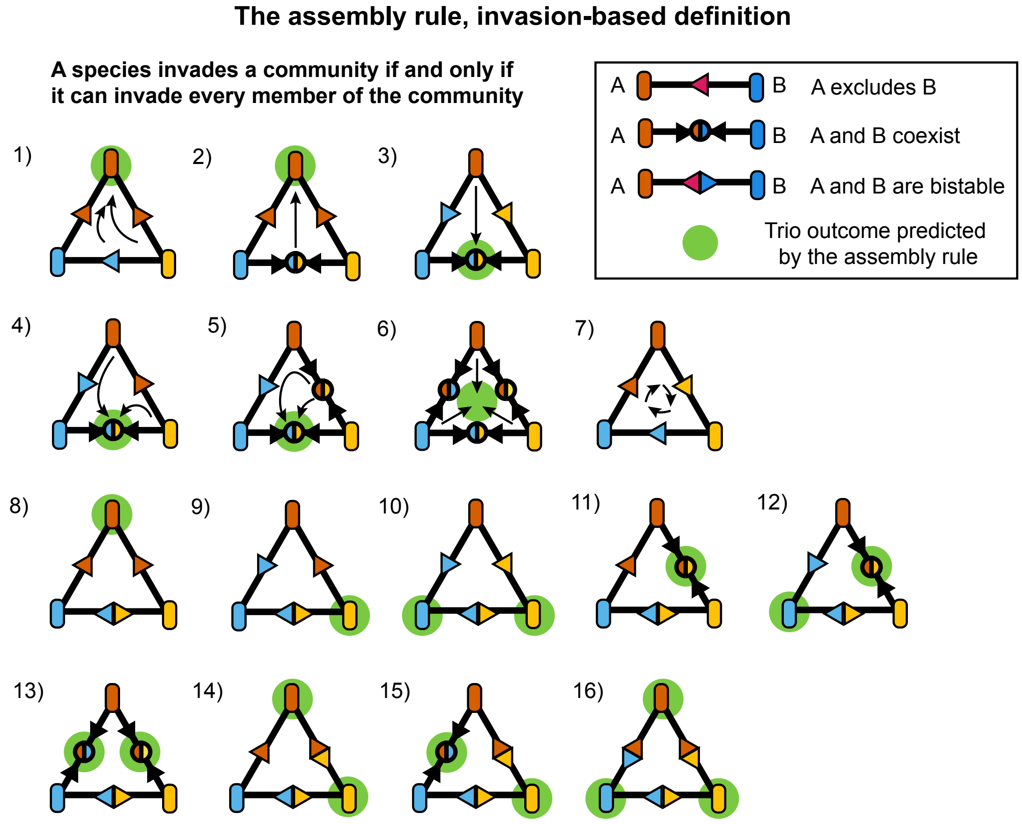


***Fig. S15. The assembly rule predictions can be extended to cases in which pairs display bistability.*** *Using the invasibility-based definition of the assembly rule, we may apply the prediction in the presence of bistability. Outcomes 1-7 involve zero bistable pair, as in Fig. 1A. Outcomes 8-13 contain one bistable pair. Outcomes 14-15 contain two bistable pairs. Outcome 16 when all three pairs are bistable. Multiple green filled circles for a trio imply alternative stable points; trio outcome can be one of the marked outcomes depending on initial species abundance.*

V.2. Resource competition with varying yield can lead to bistable pairs

Let us recall the graphical approach discussed in the main text. When biomass yields are uniform, a species that grows faster on resource 1 than resource 2 will also consume more of resource 1 than resource 2. This drives the coexistence of pairs when two species grow faster than each other on different resources. For example, let us assume species A grows faster than species B on resource 1 and more slowly than species B on resource 2. Then an increased abundance of species A will increase the consumption of resource 1, and thereby lower the concentration of resource 1, modifying the environment in favor of species B.
When biomass yields are not uniform, a species may consume more of resource on which it grows more slowly. This 'growth-yield tradeoff' scenario may lead to bistability. As an example, let's revisit the above A-B pair but now assuming that A consumes more of resource 2. Then, an increased abundance of species A will lower the concentration of resource 2 and modify the environment in favor of itself. Such positive feedback can destabilize the fixed point, which is the intersection of ZNGIs, and make the pair mutually uninvadable.

This comparison between coexistence and bistability becomes more apparent with graphical analysis because the stability of pairwise coexistence is determined by the ordering between consumption vectors of the two species. This idea is illustrated in the figure below.

**A.** **Set of possible pairwise outcomes depends on species' growth parameters**

| | No tradeoff | Growth tradeoff | Growth-yield tradeoff |
|---|---|---|---|
| Growth rate | $r_{A1} > r_{B1}$<br>$r_{A2} > r_{B2}$ | $r_{A1} > r_{B1}$<br>$r_{A2} < r_{B2}$ | $r_{A1} > r_{B1}$<br>$r_{A2} < r_{B2}$ |
| Biomass yied | - | $\frac{Y_{A1}Y_{B2}}{Y_{A2}Y_{B1}} < \frac{r_{A1}r_{B2}}{r_{A2}r_{B1}}$ | $\frac{Y_{A1}Y_{B2}}{Y_{A2}Y_{B1}} > \frac{r_{A1}r_{B2}}{r_{A2}r_{B1}}$ |
| Possible pairwise outcomes | Exclusion (A wins B) | Exclusions, Coexistence | Exclusions, Bistability |

**B.** **Competition outcome for each species pair depends on resources supply**

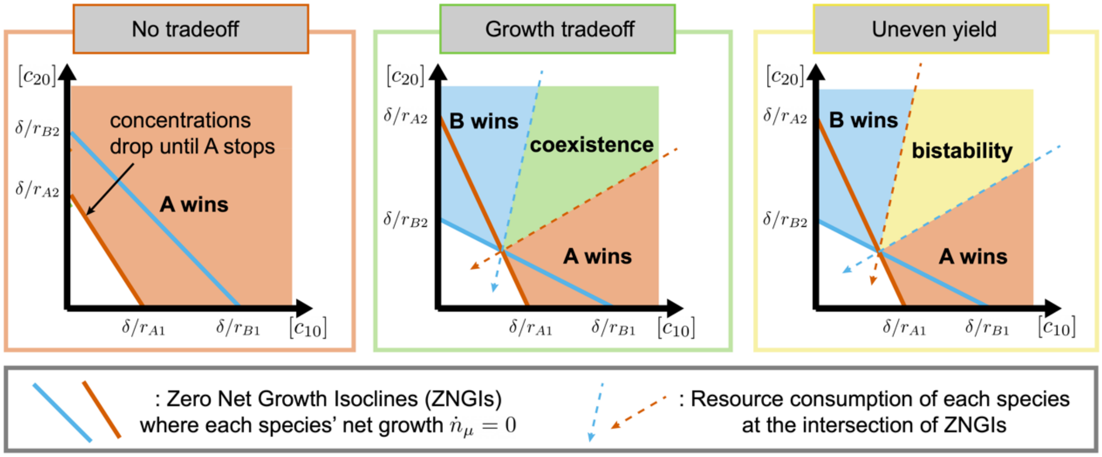


V.3. Resource competition with bistable pairs often follow the assembly rule

To study how presence of bistability affect the prediction accuracy of the assembly rule, we ran resource competition simulations of three species on three resources with varying yields. In simulations of each trio community, both per-capita per-resource growth rates $r_{\mu i}$ and inverse biomass yields $1/Y_{\mu i}$ are drawn from uniform distribution between 0 and 1 and then, for each trio community, normalized to $\sum_\mu \sum_i r_{\mu i} = \sum_\mu \sum_i \frac{1}{Y_{\mu i}} = 1$. Resource supply is set as $c_i^s = 1$ for all three resources. Then *in silico* competitions ran with 10 different initial abundances ((0.01,0.99) and (0.99,0.01) for each pair, (0.99,0.05,0.05), (0.05,0.99,0.05), (0.05,0.05,0.99), and (0.33,0.33,0.33) for trio) as outcomes may depend on initial community composition. Final community composition for each competition is solved with ODE solver. Out of 3332 such simulations with three species and three resources, we got 194 cases with one or more bistable pairs.

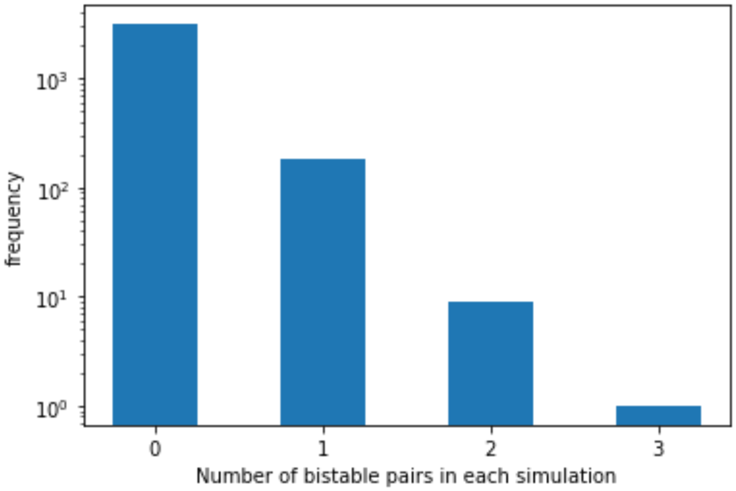


Focusing on the simulated community assemblies with one or more bistable pairs, we checked how the assembly rule prediction worked. Out of 194 simulated trios, only 3 resulted in trio outcome that is not predicted by the assembly rule. There were also 17 simulations in which one possible trio outcome (out of multiple bistable outcomes) was not observed. Except those, all simulations agreed with the assembly rule and exhibited all the predicted trio outcomes.

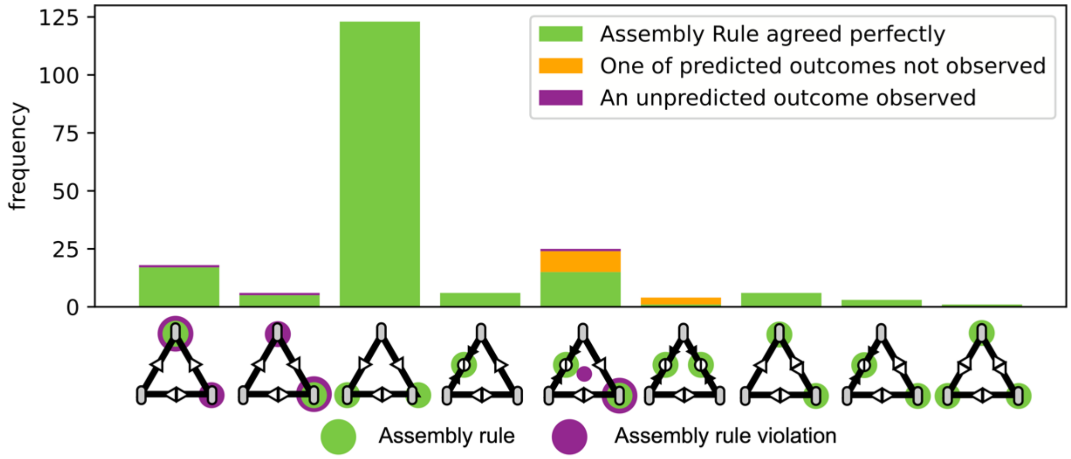


Overall, we find that community assembly with bistable pairs is well predicted by the assembly rule.
